# Extracellular matrix stiffness and TGFβ2 regulate YAP/TAZ activity in human trabecular meshwork cells

**DOI:** 10.1101/2021.09.29.462394

**Authors:** Haiyan Li, VijayKrishna Raghunathan, W. Daniel Stamer, Preethi S. Ganapathy, Samuel Herberg

## Abstract

Primary open-angle glaucoma progression is associated with increased human trabecular meshwork (HTM) stiffness and elevated transforming growth factor beta 2 (TGFβ2) levels in aqueous humor. Increased transcriptional activity of Yes-associated protein (YAP) and transcriptional coactivator with PDZ-binding motif (TAZ), central players in mechanotransduction, are implicated in glaucomatous HTM (GTM) cell dysfunction. Yet, the detailed mechanisms underlying YAP/TAZ modulation in HTM cells in response to alterations in extracellular matrix (ECM) stiffness and TGFβ2 levels are not well understood. Using biomimetic ECM hydrogels with tunable stiffness, here we show that increased ECM stiffness elevates YAP/TAZ transcriptional activity potentially through modulating focal adhesions and cytoskeletal rearrangement. Furthermore, TGFβ2 increased YAP/TAZ nuclear localization in both normal and glaucomatous HTM cells, which was prevented by inhibiting extracellular-signal-regulated kinase and Rho-associated kinase signaling pathways. Filamentous (F)-actin depolymerization reversed TGFβ2-induced YAP/TAZ nuclear localization. YAP/TAZ depletion using siRNA or verteporfin decreased focal adhesions, ECM remodeling and cell contractile properties. Similarly, YAP/TAZ inactivation with verteporfin partially blocked TGFβ2-induced HTM/GTM hydrogel contraction and stiffening. Collectively, our data provide evidence for a pathologic role of aberrant YAP/TAZ signaling in glaucomatous HTM cell dysfunction, and may help inform strategies for the development of novel multifactorial approaches to prevent progressive ocular hypertension in glaucoma.

## 1. Introduction

Primary open-angle glaucoma (POAG) is a leading cause of irreversible vision loss worldwide, and elevated intraocular pressure (IOP) is the primary modifiable risk factor (Quigley, 1993; Quigley and Broman, 2006; Kwon et al., 2009; Tham et al., 2014; Tamm et al., 2015). Elevated IOP results from increased resistance to aqueous humor outflow in the juxtacanalicular region of the conventional outflow pathway where the trabecular meshwork (TM) and Schlemm’s canal inner wall cells interact (Brubaker, 1991). Human trabecular meshwork (HTM) cells within the JCT tissue are surrounded by a complex extracellular matrix (ECM), which is primarily composed of non-fibrillar and fibrillar collagens, elastic fibrils, proteoglycans and the glycosaminoglycan hyaluronic acid (Acott and Kelley, 2008; Tamm, 2009; Hann and Fautsch, 2011; Keller and Acott, 2013; Abu-Hassan et al., 2014). In addition to providing structural support, the ECM imparts biochemical signals (e.g., hormones, growth factors and diffusible morphogens) and mechanical cues (e.g., matrix stiffness; tensile, compressive and shear forces; topographical strain) to regulate cell behaviors (Frantz et al., 2010). These external biophysical cues are translated into internal biochemical signaling cascades through a process known as mechanotransduction, and cells play an active role in remodeling their matrix to promote mechanical homeostasis and maintain tissue-level functionality (Lampi and Reinhart-King, 2018).

HTM cells are exposed to a variety of biophysical cues through cell-cell and cell-ECM interactions, and fluctuations in IOP. In POAG, impaired HTM cell function (i.e., remodeling of cell cytoskeleton, increased cell stiffness) and increased ECM deposition contribute to HTM stiffening (Schlunck et al., 2008; Han et al., 2011; McKee et al., 2011; Wang et al., 2017b; Li et al., 2021a). HTM stiffening affects HTM cell function and IOP in a feed-forward cycle characterized by dynamic reciprocity. HTM stiffening in POAG indicates that a biophysical component likely contributes to IOP regulation. Therefore, targeting mechanotransduction pathways in the HTM to interrupt this feed-forward cycle between HTM stiffening and cellular response to the stiffened ECM is emerging as a promising strategy for managing ocular hypertension.

Yes-associated protein (YAP) and transcriptional coactivator with PDZ-binding motif (TAZ, encoded by WWTR1) are central players in mechanotransduction. YAP/TAZ can be activated by increased ECM stiffness, mechanical stress and growth factors (Dupont et al., 2011; Boopathy and Hong, 2019). It has been demonstrated that certain glaucoma related molecules that influence outflow function, such as dexamethasone, lysophosphatidic acid and interleukin-6, mediate the expression of YAP and TAZ in HTM cells (Honjo et al., 2018; Peng et al., 2018; Yemanyi and Raghunathan, 2020). YAP was found to be reduced in HTM cells incubated on stiff two-dimensional (2D) polyacrylamide hydrogels, while TAZ levels were increased (Raghunathan et al., 2013). Other research showed that both YAP and TAZ were significantly higher in HTM cells on stiffer polyacrylamide substrates or stiffened HTM cell-derived matrices (Thomasy et al., 2013; Yemanyi et al., 2020b). These apparent contradictory findings suggest that the precise role of YAP/TAZ in HTM mechanotransduction is far from clear. The substrate and context dependency of YAP/TAZ signaling in these two studies highlights the importance of accurately mimicking the native tissue to further our understanding of HTM cell mechanobiology with focus on cell-ECM interactions. This is especially important considering a recent multi-ethnic genome wide meta-analysis report that identified *YAP1* as a potential genetic risk factor for POAG across European, Asian, and African ancestries implicating a causal relationship for outflow dysfunction (Gharahkhani et al., 2021b).

Biomaterials for cell culture applications are designed to mimic aspects of the ECM and provide researchers with methods to better understand cell behaviors. Protein-based hydrogels (i.e., water-swollen networks of polymers) are attractive biomaterials owing to their similarity to *in vivo* cellular microenvironments, biocompatibility, biodegradability and tunable mechanical properties (Drury and Mooney, 2003; Panahi and Baghban-Salehi, 2019). Recently, we developed a bioengineered hydrogel composed of HTM cells and ECM biopolymers found in the native HTM tissue, and demonstrated its viability for investigating cell-ECM interactions under normal and simulated glaucomatous conditions in both 2D and 3D conformations (Li et al., 2021a; Li et al., 2021b).

Here, we hypothesized that altered YAP/TAZ transcriptional activity drives HTM cell dysfunction under simulated glaucomatous conditions. The TM from POAG eyes is stiffer than that from healthy eyes (Last et al., 2011; Wang et al., 2017a; Vahabikashi et al., 2019); transforming growth factor beta 2 (TGFβ2), the predominant TGFβ isoform in the eye and aqueous humor, has been identified as a major contributor to the pathologic changes occurring in ocular hypertension and POAG (Granstein et al., 1990; Quigley, 1993; Inatani et al., 2001; Fuchshofer and Tamm, 2009; Agarwal et al., 2015; Kasetti et al., 2018). Using TGFβ2 as a glaucomatous stimulus and tuning the stiffness of our hydrogels, here, we investigated the effects of ECM stiffening and TGFβ2 on regulating YAP/TAZ transcriptional activity in HTM cells in 2D and 3D cultures. In this tissue-mimetic environment, we then investigated whether YAP/TAZ inhibition would alleviate ECM stiffness- or TGFβ2-induced HTM cell pathobiology.

## 2. Materials and Methods

### 2.1 HTM cell isolation and culture

Experiments using human donor eye tissue were approved by the SUNY Upstate Medical University Institutional Review Board (protocol #1211036), and were performed in accordance with the tenets of the Declaration of Helsinki for the use of human tissue. Primary HTM cells were isolated from healthy donor corneal rims discarded after transplant surgery as recently described (Li et al., 2021a; Li et al., 2021b), and cultured according to established protocols (Stamer et al., 1995; Keller et al., 2018). Five HTM cell strains (HTM05, HTM12, HTM14, HTM17, HTM19) were used for the experiments in this study (**Suppl. Table. 1**). All HTM cell strains were validated with dexamethasone (DEX; Fisher Scientific, Waltham, MA, USA; 100 nM) induced myocilin (MYOC) expression in more than 50% of cells by immunocytochemistry and immunoblot analyses (**Suppl. Fig. 1A,B** and (Li et al., 2021a; Li et al., 2021b)). Different combinations of 2-3 HTM cell strains were used per experiment with 3-4 replicates each, depending on cell availability, and all studies were conducted between cell passage 3-7. HTM cells were cultured in low-glucose Dulbecco’s Modified Eagle’s Medium (DMEM; Gibco; Thermo Fisher Scientific) containing 10% fetal bovine serum (FBS; Atlanta Biologicals, Flowery Branch, GA, USA) and 1% penicillin/streptomycin/glutamine (PSG; Gibco), and maintained at 37°C in a humidified atmosphere with 5% CO_2_. Fresh media was supplied every 2-3 days.

### 2.2 Glaucomatous donor history

Glaucomatous HTM cells (GTM211 and GTM1445) were used in this study. GTM211 was isolated from a 75-year-old, white female that was taking Xalatan (0.005%) in both eyes nightly for treatment of ocular hypertension. The patient was diagnosed with glaucoma, and had bilateral cataract surgery, and retina surgery. Upon receipt of donor eyes by W.D.S., outflow facility measurements during constant pressure perfusion conditions were 0.24 and 0.17 μl/min/mmHg from OD and OS eyes, respectively. GTM211 were isolated from the OS eye of the donor and characterized as previously described (Li et al., 2021a). GTM1445 has been characterized in our previous studie (Li et al., 2021a). GTM cell strain information can be found in **Suppl. Table. 1**.

### 2.3 Preparation of hydrogels

Hydrogel precursors methacrylate-conjugated bovine collagen type I (MA-COL, Advanced BioMatrix, Carlsbad, CA, USA; 3.6 mg/ml [all final concentrations]), thiol-conjugated hyaluronic acid (SH-HA, Glycosil®, Advanced BioMatrix; 0.5 mg/ml, 0.025% (w/v) photoinitiator Irgacure® 2959; Sigma-Aldrich, St. Louis, MO, USA) and in-house expressed elastin-like polypeptide (ELP, thiol via K<u>C</u>TS flanks (Li et al., 2021a); 2.5 mg/ml) were thoroughly mixed. Thirty microliters of the hydrogel solution were pipetted onto a Surfasil (Fisher Scientific) coated 12-mm round glass coverslip followed by placing a regular 12-mm round glass coverslip onto the hydrogels. Constructs were crosslinked by exposure to UV light (OmniCure S1500 UV Spot Curing System; Excelitas Technologies, Mississauga, Ontario, Canada) at 320-500 nm, 2.2 W/cm^2^ for 5 s, as previously described (Li et al., 2021a). The hydrogel-adhered coverslips were removed with fine-tipped tweezers and placed in 24-well culture plates (Corning; Thermo Fisher Scientific). Hydrogel stiffening was achieved by soaking the hydrogels in 0.1% (w/v) riboflavin (RF; Sigma) for 5 min, followed by low-intensity secondary UV crosslinking (bandpass filter: 405-500 nm, 5.4 mW/cm^2^) for 5 min.

### 2.4 HTM cell treatments

HTM/GTM cells were seeded at 2 × 10^4^ cells/cm^2^ on premade hydrogels/coverslips/tissue culture plates, and cultured in DMEM with 10% FBS and 1% PSG for 1 or 2 days. Then, HTM cells were cultured in serum-free DMEM with 1% PSG and subjected to the different treatments for 3 d: TGFβ2 (2.5 ng/ml; R&D Systems, Minneapolis, MN, USA), the ERK inhibitor U0126 (10 μM; Promega, Madison, WI, USA), the Rho-associated kinase (ROCK) inhibitor Y27632 (10 μM; Sigma-Aldrich), the actin de-polymerizer latrunculin B (10 μM; Tocris Bioscience; Thermo Fisher Scientific), or the YAP inhibitor verteporfin (0.5 μM; Sigma). The monolayer HTM cells were processed for immunoblot, qRT-PCR and immunocytochemistry analyses.

### 2.5 Immunoblot analysis

Protein was extracted from cells using lysis buffer (CelLytic™ M, Sigma-Aldrich) supplemented with Halt™ protease/phosphatase inhibitor cocktail (Thermo Fisher Scientific). Equal protein amounts (10 μg), determined by standard bicinchoninic acid assay (Pierce; Thermo Fisher Scientific), in 4× loading buffer (Invitrogen; Thermo Fisher Scientific) with 5% beta-mercaptoethanol (Fisher Scientific) were boiled for 5 min and subjected to SDS-PAGE using NuPAGE™ 4-12% Bis-Tris Gels (Invitrogen; Thermo Fisher Scientific) at 120V for 80 min and transferred to 0.45 μm PVDF membranes (Sigma; Thermo Fisher Scientific). Membranes were blocked with 5% bovine serum albumin (Thermo Fisher Scientific) in tris-buffered saline with 0.2% Tween®20 (Thermo Fisher Scientific), and probed with various primary antibodies followed by incubation with HRP-conjugated secondary antibodies or fluorescent secondary antibodies (LI-COR, Lincoln, NE, USA). Bound antibodies were visualized with the enhanced chemiluminescent detection system (Pierce) on autoradiography film (Thermo Fisher Scientific) or Odyssey® CLx imager (LI-COR). Densitometry was performed using the open-source National Institutes of Health software platform, FIJI (Schindelin et al., 2012) or Image Studio™ Lite (LI-COR); data were normalized to GAPDH. A list of all of antibodies used in this study, including their working dilutions, can be found in **Supplementary Table 2**.

### 2.6 Immunocytochemistry analysis

HTM cells in presence of the different treatments were fixed with 4% paraformaldehyde (Thermo Fisher Scientific) at room temperature for 20 min, permeabilized with 0.5% Triton™ X-100 (Thermo Fisher Scientific), blocked with blocking buffer (BioGeneX), and incubated with primary antibodies, followed by incubation with fluorescent secondary antibodies; nuclei were counterstained with 4′,6′-diamidino-2-phenylindole (DAPI; Abcam). Similarly, cells were stained with Phalloidin-iFluor 488 or 594 (Abcam)/DAPI according to the manufacturer’s instructions. Coverslips were mounted with ProLong™ Gold Antifade (Invitrogen) on Superfrost™ microscope slides (Fisher Scientific), and fluorescent images were acquired with an Eclipse N*i* microscope (Nikon Instruments, Melville, NY, USA) or a Zeiss LSM 780 confocal microscope (Zeiss, Germany). Fluorescent signal intensity was measured using FIJI software. A list of all of antibodies used in this study, including their working dilutions, can be found in **Supplementary Table 2**.

### 2.7 Image analysis

All image analysis was performed using FIJI software. Briefly, the cytoplasmic YAP intensity was measured by subtracting the overlapping nuclear (DAPI) intensity from the total YAP intensity. The nuclear YAP intensity was recorded as the proportion of total YAP intensity that overlapped with the nucleus (DAPI). YAP/TAZ nuclear/cytoplasmic (N/C) ratio was calculated as follows: N/C ratio = (nuclear YAP signal/area of nucleus)/(cytoplasmic signal/area of cytoplasm). Fluorescence intensity of F-actin, αSMA, FN, and p-MLC were measured in at least 20 images from 2 HTM cell strains with 3 replicates per HTM cell strain with image background subtraction using FIJI software. For number of vinculin puncta quantification, the 3D Object Counter plugin in FIJI software was used to threshold images and quantify puncta/0.05 mm^2^ (above 2 μm in size). Nuclear area was also measured using FIJI software. At least 100 nuclei were analyzed from 2 HTM cell strains with 3 replicates per HTM cell strain.

### 2.8 Quantitative reverse transcription-polymerase chain reaction (qRT-PCR) analysis

Total RNA was extracted from HTM cells using PureLink RNA Mini Kit (Invitrogen). RNA concentration was determined with a NanoDrop spectrophotometer (Thermo Fisher Scientific). RNA was reverse transcribed using iScript™ cDNA Synthesis Kit (BioRad, Hercules, CA, USA). One hundred nanograms of cDNA were amplified in duplicates in each 40-cycle reaction using a CFX 384 Real Time PCR System (BioRad) with annealing temperature set at 60ºC, Power SYBR™ Green PCR Master Mix (Thermo Fisher Scientific), and custom-designed qRT-PCR primers. Transcript levels were normalized to GAPDH, and mRNA fold-changes calculated relative to vehicle controls using the comparative C_T_ method (Schmittgen and Livak, 2008). A list of all of primers used in this study can be found in **Supplementary Table 3**.

### 2.9 Active TGFβ2 enzyme-linked immunosorbent assay (ELISA)

TGFβ2 levels were quantified using the Quantikine ELISA kit (R&D Systems). HTM and GTM cells were seeded at 2 × 10^4^ cells/cm^2^ on tissue culture plates, and cultured in DMEM with 10% FBS and 1% PSG for 1 or 2 days to grow to 80 – 90 % confluence. Then, HTM and GTM cells were cultured in 1 ml serum-free DMEM with 1% PSG for 3 d. After 3 days, the cell culture supernatants were collected and centrifuged at 1000 g for 10 min. The supernatants were used for ELISA according to the manufacturer’s instructions.

### 2.10 siRNA transfection

HTM cells were depleted of YAP and TAZ using siRNA-loaded lipofectamine RNAimax (Invitrogen) according to the manufacturer’s instructions. In brief, HTM cells were seeded at 2 × 10^4^ cells/cm^2^ on RF double-crosslinked hydrogels in DMEM with 10% FBS and 1% PSG. The following day, the cell culture medium was changed to antibiotic-free and serum-free DMEM and the samples were kept in culture for 24 h followed by transfection. Transfection was performed using a final concentration 3% (v/v) lipofectamine RNAimax with 150 nM RNAi duplexes (custom oligonucleotides; Dharmacon). Transfected HTM cells were used 48 h after transfection. ON-TARGET plus nontargeting siRNA were obtained from Dharmacon. Custom siRNA were based on sequences previously described (Dupont et al., 2011): YAP, sense, 5’-GACAUCUUCUGGUCAGAGA-3’, and YAP, anti-sense, 5’-UCUCUGACCAGAAGAUGUC-3’; TAZ, sense, 5’-ACGUUGACUUAGGAACUUU-3’, and TAZ, anti-sense, 5’-AAAGUUCCUAAGUCAACGU-3’.

### 2.11 HTM hydrogel contraction analysis

HTM cell-laden hydrogels were prepared by mixing HTM cells (1.0 × 10^6^ cells/ml) with MA-COL (3.6 mg/ml [all final concentrations]), SH-HA (0.5 mg/ml, 0.025% (w/v) photoinitiator) and ELP (2.5 mg/ml) on ice, followed by pipetting 10 μl droplets of the HTM cell-laden hydrogel precursor solution onto polydimethylsiloxane (PDMS; Sylgard 184; Dow Corning) coated 24-well culture plates. Constructs were crosslinked as described above (320-500 nm, 2.2 W/cm^2^, 5 s). HTM cell-laden hydrogels were cultured in DMEM with 10% FBS and 1% PSG in presence of the different treatments. Longitudinal brightfield images were acquired at 0 d and 5 d with an Eclipse T*i* microscope (Nikon). Construct area from N = 8-11 hydrogels per group from 2 HTM/GTM cell strains with 3-4 biological replicates per HTM/GTM cell strain was measured using FIJI software and normalized to 0 d followed by normalization to controls.

### 2.12 HTM hydrogel cell proliferation analysis

Cell proliferation was measured with the CellTiter 96® Aqueous Non-Radioactive Cell Proliferation Assay (Promega) following the manufacturer’s protocol. HTM hydrogels cultured in DMEM with 10% FBS and 1% PSG in presence of the different treatments for 5 d were incubated with the staining solution (38 μl MTS, 2 μl PMS solution, 200 μl DMEM) at 37ºC for 1.5 h. Absorbance at 490 nm was recorded using a spectrophotometer plate reader (BioTEK, Winooski, VT, USA). Blank-subtracted absorbance values served as a direct measure of HTM cell proliferation from N = 6-8 hydrogels per group from 2 HTM/GTM cell strains with 3-4 biological replicates per HTM/GTM cell strain.

### 2.13 HTM hydrogel rheology analysis

Fifty microliters of acellular hydrogel precursor solutions were pipetted into custom 8×1-mm PDMS molds. Similarly, 250 μl of HTM cell-laden hydrogel precursor solutions were pipetted into 16×1-mm PDMS molds. All samples were UV crosslinked and equilibrated as described above. Acellular hydrogels were measured at 0 d. HTM hydrogels, cultured in DMEM with 10% FBS and 1% PSG in presence of the different treatments, were measured at 5 d; samples were cut to size using an 8-mm diameter tissue punch. A Kinexus rheometer (Malvern Panalytical, Westborough, MA, USA) fitted with an 8-mm diameter parallel plate was used to measure hydrogel viscoelasticity. To ensure standard conditions across all experiments (N = 3 per group), the geometry was lowered into the hydrogels until a calibration normal force of 0.02 N was achieved. Subsequently, an oscillatory shear-strain sweep test (0.1-60%, 1.0 Hz, 25°C) was applied to determine storage modulus (G’) and loss modulus (G”) in the linear region. Elastic modulus was calculated with E = 2 * (1 + v) * G′, where a Poisson’s ratio (v) of 0.5 for the ECM hydrogels was assumed (Timothy P. Lodge, 2020).

### 2.14 HTM hydrogel atomic force microscopy (AFM) analysis

Thirty microliters of HTM cell-laden hydrogel solution were pipetted onto a Surfasil (Fisher Scientific) coated 12-mm round glass coverslip followed by placing a regular 12-mm round glass coverslip onto the hydrogels. Constructs were crosslinked by exposure to UV light at 320-500 nm, 2.2 W/cm^2^ for 5 s, and cultured in DMEM with 10% FBS and 1% PSG for 7 days. Samples were shipped overnight to UHCO where they were locally masked. Elastic modulus of HTM cell-laden hydrogels were obtained by atomic force microscopy (AFM) in fluid in contact mode as described previously (Raghunathan et al., 2018). Briefly, force vs. indentation curves were obtained on a Bruker BioScope Resolve (Bruker nanoSurfaces, Santa Barbara, CA) AFM employing a PNP-TR cantilever (NanoAndMore, Watsonville, CA) whose pyramidal tip was modified with a borosilicate bead (nominal diameter 5 μm), nominal spring constant of 0.32 N/m assuming sample Poisson’s ratio of 0.5. The diameter of each sphere attached to the cantilever was qualified prior to usage. Prior to experimental samples, cantilever was calibrated on a silicon wafer in fluid. Force-distance curves were randomly obtained from 7-10 locations with 3 force curves obtained per location per sample. Force curves were analyzed using the Hertz model for spherical indenters using a custom MATLAB code (Chang et al., 2014). Curve fits were performed for at least 1 μm of indentation depth for the samples.

### 2.15 Statistical analysis

Individual sample sizes are specified in each figure caption. Comparisons between groups were assessed by unpaired *t*-test, one-way or two-way analysis of variance (ANOVA) with Tukey’s multiple comparisons *post hoc* tests, as appropriate. The significance level was set at p<0.05 or lower. GraphPad Prism software v9.2 (GraphPad Software, La Jolla, CA, USA) was used for all analyses.

## 3. Results

### 3.1 TGFβ2 and YAP/TAZ activity is upregulated in GTM cells

It has been shown that levels of TGFβ2 are elevated in eyes of glaucomatous patients compared to age-matched normal eyes (Inatani et al., 2001; Picht et al., 2001; Ochiai and Ochiai, 2002; Agarwal et al., 2015). We confirmed that GTM cells isolated from donor eyes with POAG history secreted significantly more active TGFβ2 protein by ∼2.44-fold (**Fig. 1A**) compared to normal HTM cells, which was consistent with increased mRNA levels (**Suppl. Fig. 2A**).

**Fig. 1.**
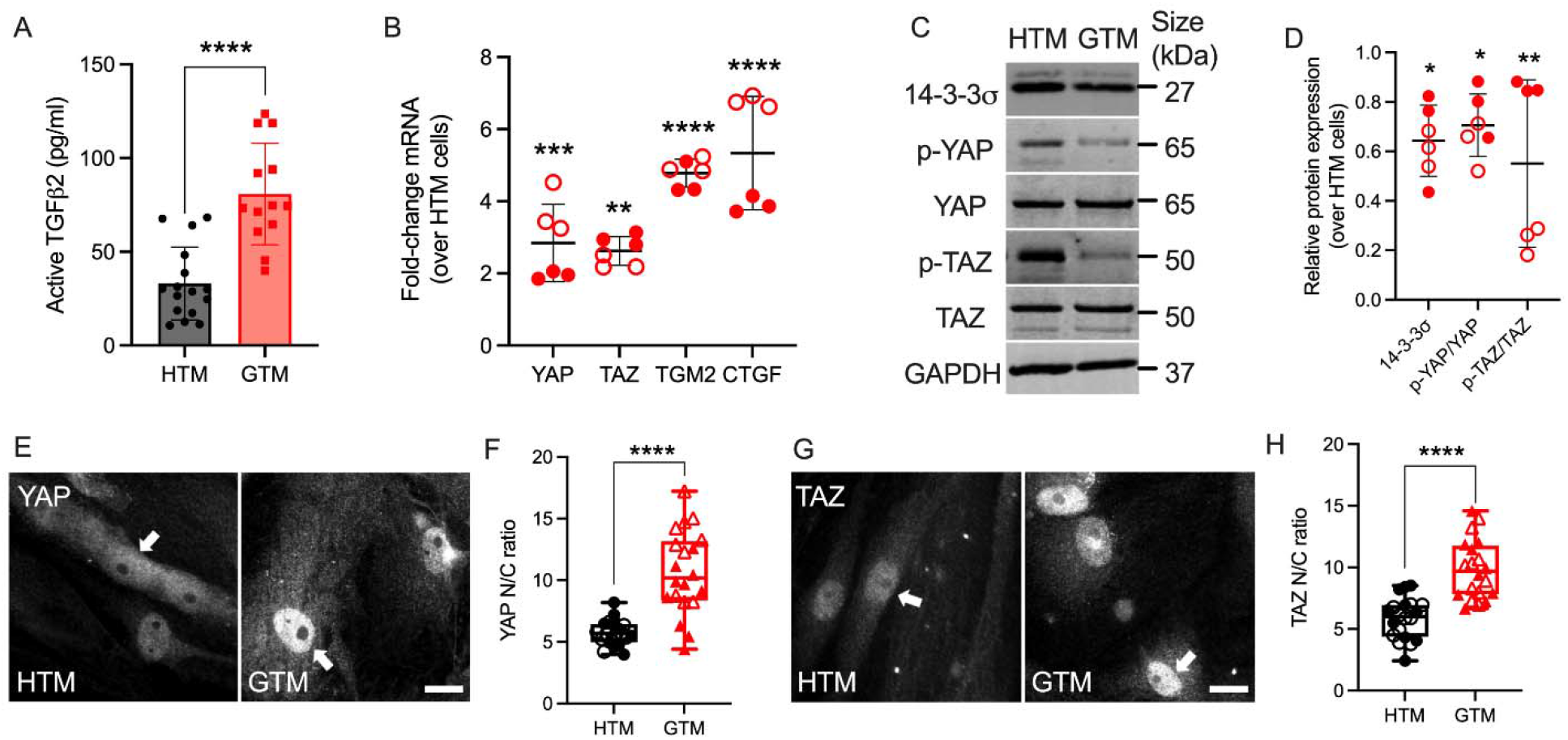
Upregulated active TGFβ2 and YAP/TAZ nuclear localization in GTM cells compared to normal HTM cells. (A) Active TGFβ2 levels were quantified by ELISA (N = 16 replicates from 3 HTM cell strains, N = 13 replicates from 2 GTM cell strains). (B) mRNA fold-change of YAP, TAZ, TGM2 and CTGF by qRT-PCR (N = 6 replicates from 2 HTM/GTM cell strains). (C) Immunoblot of 14-3-3σ, p-YAP, p-TAZ, total YAP and total TAZ. (D) Immunoblot analysis of 14-3-3σ, p-YAP/YAP and p-TAZ/TAZ (N = 6 replicates from 2 HTM/GTM cell strains). (E and G) Representative fluorescence micrographs of YAP/TAZ in HTM and GTM cells (YAP/TAZ = grey). Scale bar, 20 μm; arrows indicate YAP/TAZ nuclear localization. (F and H) Analysis of YAP/TAZ nuclear/cytoplasmic ratio (N = 20 images from 2 HTM/GTM cell strains with 3 replicates per cell strain). Open and closed symbols represent different cell strains. In A, B and D, the bars or lines and error bars indicate Mean ± SD; In F and H, the box and whisker plots represent median values (horizontal bars), 25th to 75th percentiles (box edges) and minimum to maximum values (whiskers), with all points plotted. Significance was determined by unpaired t-test (A, F and H) and two-way ANOVA using multiple comparisons tests (D) (*p < 0.05; **p < 0.01; ***p < 0.001; ****p < 0.0001).

To investigate YAP and TAZ transcriptional activity under normal and glaucomatous conditions, the expression of YAP/TAZ and relevant select downstream targets at mRNA and protein levels were evaluated in normal HTM and GTM cells, which were plated with similar cell density. The functions of YAP/TAZ depend on their spatial localization within the cellular nucleus or cytoplasm. When localized to the nucleus, YAP/TAZ interact with TEAD transcription factors to drive the expression of certain proteins, such as transglutaminase-2 (TGM2) and connective tissue growth factor (CTGF), cysteine-rich angiogenic inducer 61 (CYR61) and ankyrin repeat domain 1 (ANKRD1) (Low et al., 2014). When localized in the cytoplasm, YAP/TAZ can be phosphorylated at Ser 127 by the large tumor suppressor (LATS) 1/2 kinase; phosphorylation of YAP/TAZ either primes to binding with 14-3-3σ leading to cytoplasmic sequestration of YAP/TAZ or ubiquitin-mediated protein degradation (Zhao et al., 2007; Lei et al., 2008; Dupont et al., 2011). We demonstrated that mRNA levels of YAP and TAZ were both significantly upregulated in GTM cells compared to normal HTM cells (**Fig. 1B**). TGM2 and CTGF have been shown to play a role in HTM cell pathobiology in glaucoma (Tovar-Vidales et al., 2008; Junglas et al., 2012); here we showed that mRNA of TGM2, CTGF and ANKRD1 were also significantly upregulated in GTM cells vs. normal HTM cells, whereas no significant difference was observed for CYR61 (**Fig. 1B; Suppl. Fig. 2B,C**). GTM cells showed significantly lower p-YAP and p-TAZ vs. normal HTM cells, while total YAP and TAZ expression were similar to HTM cells, leading to decreased p-YAP/YAP and p-TAZ/TAZ ratios. Consistent with the ratio of phosphorylated-to-total proteins, 14-3-3σ expression in GTM cells was significantly decreased compared to normal HTM cells (**Fig. 1C,D**). Besides, GTM cells exhibited significantly increased YAP/TAZ nuclear-to-cytoplasmic (N/C) ratio and TGM2 expression compared to normal HTM cells (**Fig. 1E-H; Suppl. Fig. 3**). Together, these data demonstrate that levels of active TGFβ2 as well as YAP and TAZ transcriptional activity are elevated in GTM cells isolated from patients with glaucoma compared to normal HTM cells.

### 3.2 ECM stiffening increases YAP/TAZ activity in HTM cells via modulating focal adhesions and cytoskeletal rearrangement

Atomic force microscopy (AFM) analyses showed that the TM from POAG eyes is ∼1.5-5-fold stiffer compared to that from healthy eyes (Wang et al., 2017a; Wang et al., 2017b; Vahabikashi et al., 2019). To that end, we recently showed that DEX treated HTM cell-laden hydrogels are ∼2-fold stiffer compared to controls (Li et al., 2021a). Here, we demonstrated that TGFβ2-treated HTM cell encapsulated hydrogels were also ∼2-fold stiffer compared to controls using AFM and rheology (**Fig. 2A, Suppl. Fig. 4**). To mimic the stiffness difference between glaucomatous and healthy HTM tissue, we utilized riboflavin (RF)-mediated secondary UV crosslinking of collagen fibrils, which stiffened the hydrogels by ∼2-fold (**Fig. 2B**).

**Fig. 2.**
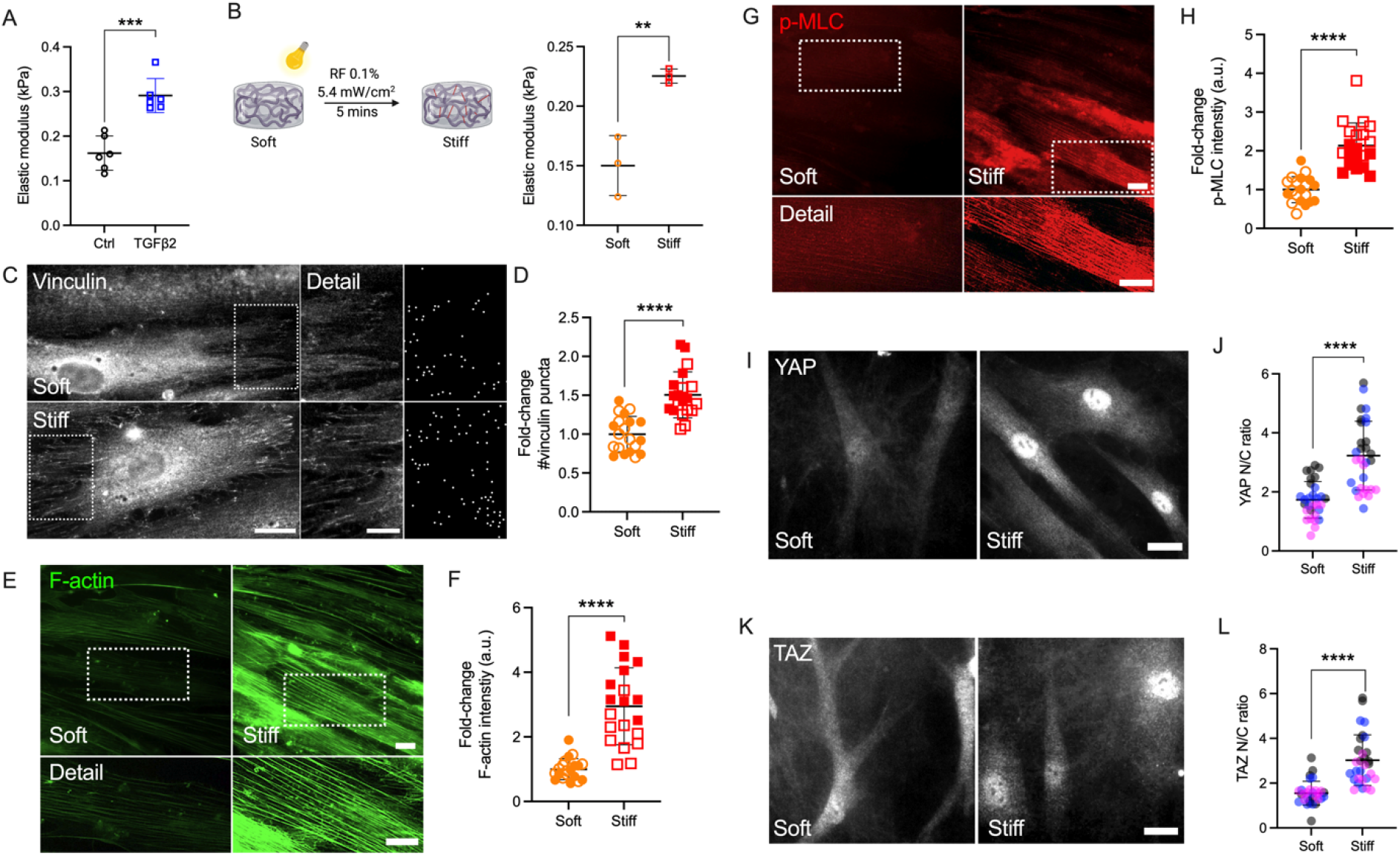
Stiffened ECM hydrogels elevate YAP/TAZ activity in HTM cells via modulating FA and cytoskeletal rearrangement. (A) Elastic modulus of HTM cell-encapsulated hydrogels subjected to control and TGFβ2 (2.5 ng/mL) measured by AFM (N = 6 replicates/group). (B) Schematic of riboflavin (RF)-mediated double photo-crosslinking to stiffen ECM hydrogels, and elastic modulus of the hydrogels (N = 3 replicates/group). (C) Representative fluorescence micrographs and FIJI mask images of vinculin in HTM cells on soft and stiff hydrogels (dashed box shows region of interest in higher magnification image; vinculin = grey). Scale bar, 20 μm (left) and 5 μm (right). (D) Analysis of number of vinculin puncta (N = 20 images from 2 HTM cell strains with 3 replicates per cell strain). (E and G) Representative fluorescence micrographs of F-actin and p-MLC in HTM cells on soft and stiff hydrogels (dashed box shows region of interest in higher magnification image. F-actin = green; p-MLC =red). Scale bar, 20 μm. (F and H) Analysis of F-actin and p-MLC intensity (N = 20 images per group from 2 HTM cell strains with 3 replicates per HTM cell strain). (I and K) Representative fluorescence micrographs of YAP/TAZ in HTM cells on soft and stiff hydrogels (YAP/TAZ = grey). Scale bar, 20 μm. (J and L) Analysis of YAP/TAZ nuclear/cytoplasmic ratio (N = 30 images from 2 HTM cell strains with 3-6 biological replicates per cell strain). Open and closed symbols, or symbols with different colors represent different cell strains. The bars and error bars indicate Mean ± SD; Significance was determined by unpaired t-test (**p < 0.01; ***p < 0.001; ****p < 0.0001).

Adherent cells are connected to the ECM through transmembrane receptor integrins and focal adhesion (FA) proteins such as vinculin (Mohammed et al., 2019). HTM cells on stiff glass exhibited strong vinculin FA staining (**Suppl. 5A**), while cells on soft hydrogels showed qualitatively fewer vinculin puncta compared to cells on glass (**Fig. 2C**). The stiffened ECM hydrogels significantly increased number and size of vinculin puncta in HTM cells compared to HTM cells on the soft hydrogel substrates (**Fig. 2C,D; Suppl. 5C**). Consistent with our observation on vinculin FA, HTM cells on glass and stiff hydrogels exhibited bigger nuclei compared to cells on the soft hydrogels (glass: ∼1.56-fold; stiff hydrogels: ∼1.26-fold) (**Suppl. Fig. 5B**). We also found that the stiffened ECM hydrogels significantly increased filamentous (F)-actin, phospho-myosin light chain (p-MLC) and α-smooth muscle actin (αSMA) levels in HTM cells compared to cells on the soft matrix (**Fig. 2E-H, Suppl. 5D,E**). Importantly, we observed that nuclear localization of YAP/TAZ, TGM2 expression, and fibronectin (FN) remodeling in HTM cells were upregulated by the stiffened ECM hydrogels compared to cells on the soft hydrogels (**Fig. 2I-L; Suppl. Fig. 5F-I**).

Together, these results suggest that the stiffened ECM hydrogels induce a bigger nuclear size in HTM cells on stiff matrix compared to cells on soft matrix. The stiffened hydrogels induce FA and actomyosin cytoskeletal rearrangement, correlating with elevated YAP/TAZ transcriptional activity in HTM cells.

### 3.3 TGFβ2 regulates YAP/TAZ activity via ERK and ROCK signaling pathways

TGFβ2 has been shown to activate canonical Smad (Smad2/3) and diverse non-Smad signaling pathways including extracellular-signal-regulated kinase (ERK), c-Jun N-terminal kinases, P38 kinases and Rho-associated kinase (ROCK) (Zhang, 2009; Prendes et al., 2013; Montecchi-Palmer et al., 2017; Ma et al., 2020). We recently demonstrated that TGFβ2 increased HTM cell contractility via ERK and ROCK signaling pathways by differentially regulating F-actin, αSMA, FN, and p-MLC in HTM cells (Li et al., 2021b). Here, we investigated whether YAP signaling in HTM cells in response to TGFβ2 induction requires ERK or ROCK. HTM or GTM cells were seeded on top of soft hydrogels treated with TGFβ2 ± U0126 (ERK inhibitor) or Y27632 (ROCK inhibitor). Consistent with our observation for cells seeded on glass (**Fig. 1**), we found significantly higher levels of nuclear YAP/TAZ and TGM2 expression in GTM cells compared to normal HTM cells on the soft ECM hydrogels (**Fig. 3; Suppl. Fig. 6**). TGFβ2 increased nuclear YAP in both HTM and GTM cells, which were blocked by co-treatment of U0126 or Y27632. Importantly, we found that YAP N/C ratio in GTM cells with ERK or ROCK inhibition were at similar levels as HTM controls (**Fig. 3A-D**).

**Fig. 3.**
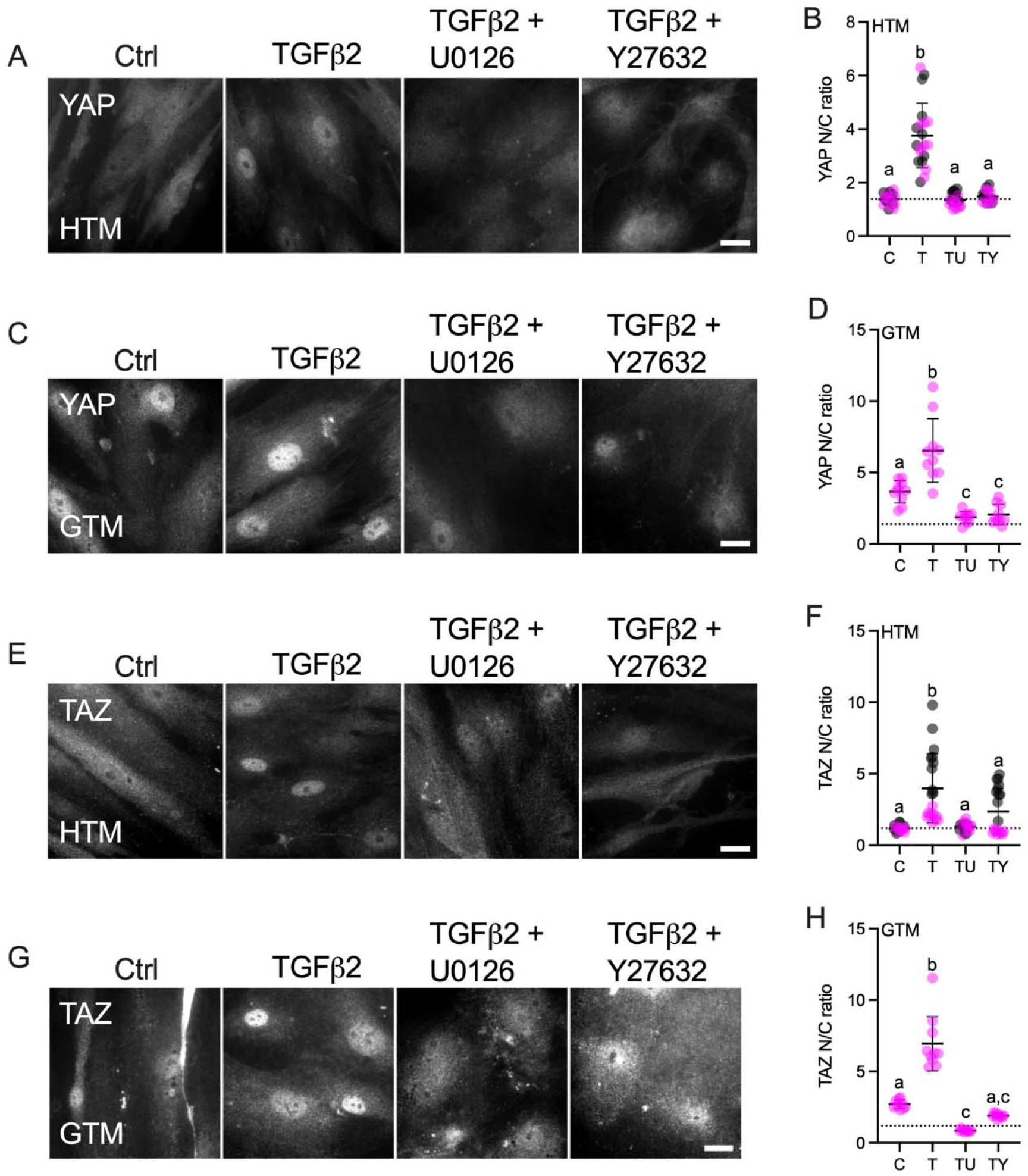
Effects of TGFβ2 in absence or presence of ERK or ROCK inhibition on nuclear YAP/TAZ localization in HTM and GTM cells. (A, C, E and G) Representative fluorescence micrographs of YAP or TAZ in HTM or GTM cells on soft hydrogels subjected to control, TGFβ2 (2.5 ng/mL), TGFβ2 + U0126 (10 μM), TGFβ2 + Y27632 (10 μM) at 3 d (YAP = grey). Scale bar, 20 μm. (B, D, F and H) Analysis of YAP or TAZ nuclear/cytoplasmic ratio (N = 20 images from 2 HTM cell strains with 3 replicates per cell strain; N = 10 images from one GTM cell strain with 3 replicates). Symbols with different colors represent different cell strains; dotted line shows HTM cells control value for reference. The bars and error bars indicate Mean ± SD. Significance was determined by one-way ANOVA using multiple comparisons tests (shared significance indicator letters represent non-significant difference (p>0.05), distinct letters represent significant difference (p<0.05)).

YAP and TAZ are generally thought to function similarly in response to mechanical and biochemical signals (Totaro et al., 2018). However, it has been shown that YAP and TAZ are distinct effectors of TGFβ1-induced myofibroblast transformation (Muppala et al., 2019). In this study, we observed that TGFβ2 significantly induced nuclear TAZ localization in HTM cells, and co-treatment of U0126 or Y27632 with TGFβ2 rescued TAZ nuclear localization to control levels (**Fig. 3E,F)**. Similarly, TGFβ2 increased TAZ N/C ratio in GTM cells, while ERK or ROCK inhibition decreased TGFβ2-induced nuclear TAZ to HTM control levels (**Fig. 3G,H**). Consistent with YAP/TAZ nuclear localization, TGM2 expression was upregulated by TGFβ2, which was significantly decreased by co-treatment of U0126 or Y27632 (**Suppl. Fig. 6**).

These data show that TGFβ2 increases nuclear YAP/TAZ and TGM2 expression in both HTM and GTM cells, which was attenuated by either U0126 or Y27632 co-treatment. Collectively, this suggests that external biophysical (stiffened ECM) and biochemical signals (increased TGFβ2) drive altered YAP and TAZ mechanotransduction in HTM cells that contributes to glaucomatous cellular dysfunction.

### 3.4 F-actin polymerization modulates YAP/TAZ activity

It has been shown that latrunculin B (Lat B), a compound that inhibits polymerization of the actin cytoskeleton, increases outflow facility and decreases IOP in human and non-human primate eyes (Peterson et al., 1999; Tian et al., 2000; Ethier et al., 2006). Likewise, Lat B has been reported to depolymerize HTM cell actin and decrease cellular stiffness via cytoskeletal relaxation (McKee et al., 2011). To investigate the effects of F-actin cytoskeletal disruption on YAP/TAZ activity, HTM cells were seeded on top of soft hydrogels, and treated with TGFβ2 for 3 d, followed by Lat B treatment for 30 mins (**Fig. 4A**). Consistent with our previous study (Li et al., 2021b), TGFβ2 significantly increased F-actin fibers, and co-treatment of Lat B with TGFβ2 potently depolymerized F-actin and decreased F-actin fiber formation (**Fig. 4B,C**) without negatively influencing cell viability (all three groups exhibited ∼8 cells/0.05mm^2^). We observed significantly more vinculin puncta and increased puncta size induced by TGFβ2, which was abolished by Lat B treatment (**Fig. 4D,E; Suppl. 7A**). Similar to before (**Fig. 3**), TGFβ2 significantly increased YAP/TAZ nuclear localization and TGM2 expression, which were restored to control levels by 30 mins of Lat B co-treatment (**Fig. 4G-J; Suppl. Fig. 7B,C**). Immunoblot analyses corroborated the immunostaining results and showed that co-treatment of TGFβ2 and Lat B increased the ratio of p-YAP to YAP and p-TAZ to TAZ, representing overall decreased nuclear YAP and TAZ. Interestingly, according to the immunoblot result, Lat B may regulate YAP activity by increasing p-YAP while maintaining similar levels of total YAP, whereas Lat B may decrease total levels of TAZ while maintaining similar levels of p-TAZ (**Fig. 4F**).

**Fig. 4.**
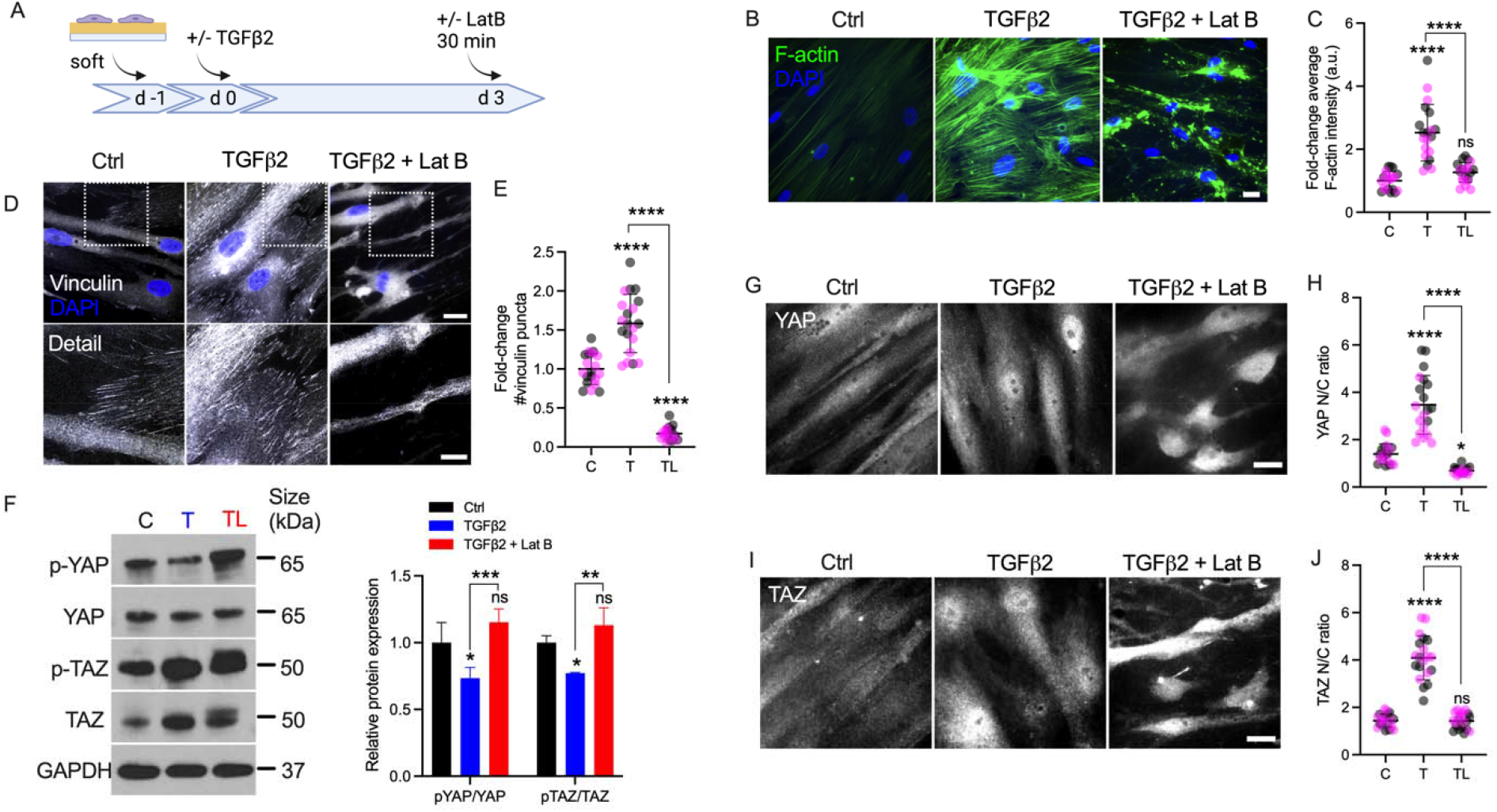
Lat B reduces YAP/TAZ activity in HTM cells. (A) Schematic showing time course of Lat B experiments with HTM cells. (B) Representative fluorescence micrographs of F-actin in HTM cells on soft hydrogels subjected to control, TGFβ2 (2.5 ng/mL), TGFβ2 + Lat B (2 μM) at 3 d (F-actin = green; DAPI = blue). Scale bar, 20 μm. (C) Quantification of F-actin intensity (N = 20 images from 2 HTM cell strains with 3 replicates per cell strain). (D) Representative fluorescence micrographs of vinculin in HTM cells on soft hydrogels subjected to the different treatments (dashed box shows region of interest in higher magnification image; vinculin = grey; DAPI = blue). Scale bar, 20 and 10 μm. (E) Analysis of number of vinculin puncta (N = 20 images from 2 HTM cell strains with 3 replicates per cell strain). (F) Immunoblot of p-YAP, total YAP, p-TAZ and total TAZ, and immunoblot analysis of p-YAP/YAP and pTAZ/TAZ (N = 3 replicates). (G and I) Representative fluorescence micrographs of YAP/TAZ in HTM cells on soft hydrogels subjected to the different treatments (YAP/TAZ = grey). Scale bar, 20 μm. (H and J) Analysis of YAP/TAZ nuclear/cytoplasmic ratio (N = 20 images from 2 HTM cell strains with 3 replicates per cell strain). Symbols with different colors represent different cell strains. The bars and error bars indicate Mean ± SD. Significance was determined by one-way (C, E, H and J) and two-way ANOVA (F) using multiple comparisons tests (*p < 0.05; **p < 0.01; ***p < 0.001; ****p < 0.0001).

In sum, these findings demonstrate that F-actin depolymerization decreases TGFβ2-induced vinculin FA, nuclear YAP/TAZ and their downstream target TGM2.

### 3.5 YAP and TAZ regulate FA formation, ECM remodeling and cell contractile properties

Our results so far suggest that POAG-related stimuli (i.e., TGFβ2 and stiffened ECM) increase YAP/TAZ activity in HTM cells. To further investigate the roles of YAP/TAZ in HTM cell function, we seeded HTM cells on stiff hydrogels to increase the baseline levels of YAP/TAZ in controls, and depleted YAP and TAZ using combined siRNA knockdown (**Fig. 5A**). Treatment reduced mRNA expression levels to 34.52% and 35.25% of siRNA controls, respectively (**Suppl. Fig. 8A**). Immunostaining showed that YAP/TAZ nuclear localization in HTM cells transfected with siYAP/TAZ were significantly decreased vs. controls despite culture in a stiffened ECM environment (**Suppl. Fig. 8B-E**). Importantly, we observed that YAP/TAZ depletion significantly reduced expression of TGM2, FN and CTGF, which may decrease HTM ECM stiffness (**Fig. 5B-E,O**).

**Fig. 5.**
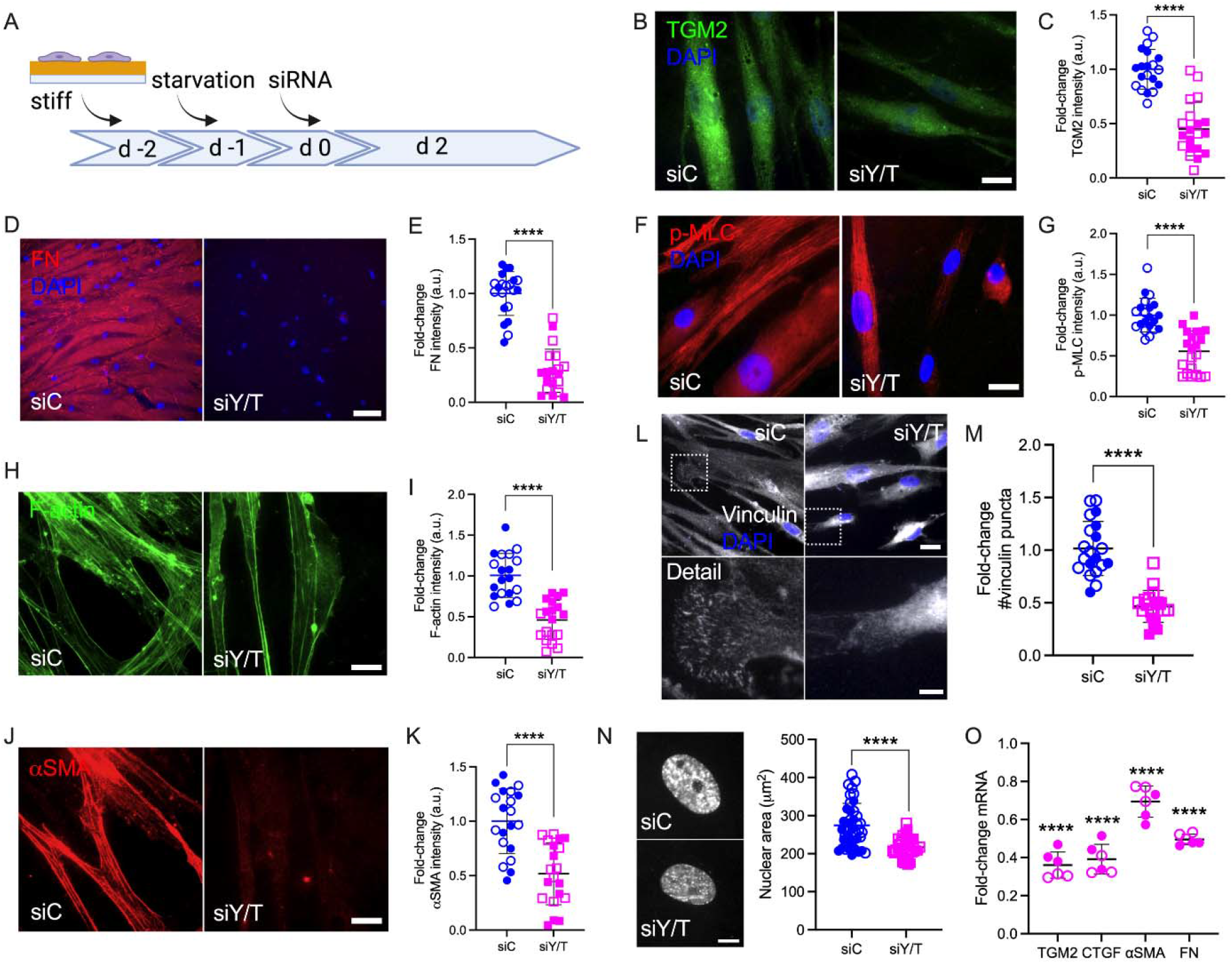
YAP/TAZ are regulators of FA formation, ECM remodeling and cell contractile properties in HTM cells. (A) Schematic showing time course of YAP/TAZ depletion using siRNA experiments with HTM cells. (B, D, F, H, J and L) Representative fluorescence micrographs of TGM2, FN, αSMA, F-actin, p-MLC and vinculin in HTM cells on stiff hydrogels subjected to siControl or siYAP/TAZ (dashed box shows region of interest in higher magnification image; TGM2/F-actin = green; FN/p-MLC/αSMA = red; vinculin = grey; DAPI = blue). Scale bar, 20 μm in B, F, H and J; 20 μm and 5 μm in L; 100 μm in D. (C, E, G, I, K and M) Analysis of TGM2, FN, αSMA, F-actin, p-MLC and number of vinculin puncta (N = 20 images from 2 HTM cell strains with 3 replicates per cell strain). (N) Representative fluorescence micrographs and analysis of nuclei of HTM cells on stiff hydrogels subjected to siControl or siYAP/TAZ (N = 40 images from 2 HTM cell strains with 6 replicates per cell strain; more than 100 nuclei were analyzed per cell strain). (O) mRNA fold-change of FN, TGM2, CTGF, αSMA in HTM cells on stiff hydrogels subjected to siControl or siYAP/TAZ by qRT-PCR. The mRNA levels were normalized to the levels of GAPDH mRNA 48h post transfection (N = 6 replicates from 2 HTM cell strain). Open and closed symbols represent different cell strains. The bars and error bars indicate Mean ± SD; Significance was determined by unpaired t-test (C, E, G, I, K, M and N) and two-way ANOVA (O) using multiple comparisons tests (**p < 0.01; ****p < 0.0001).

F-actin filaments, αSMA and p-MLC are all involved in cell contractility regulation, and we have previously demonstrated that their expression was upregulated in HTM cells under simulated glaucomatous conditions (Li et al., 2021a; Li et al., 2021b). Here, YAP/TAZ depletion significantly decreased F-actin filaments and expression of αSMA and p-MLC, suggestive of reduced HTM cell contractility (**Fig. 5F-I,O**). We also found that the expression of canonical Hippo pathway kinases LATS1, LATS2 and 14-3-3σ was not affected by siYAP/TAZ knockdown (**Suppl. Fig. 8F**). Tension generated within the actomyosin cytoskeleton is transmitted across FA to induce integrin-mediated remodeling of the ECM (Jansen et al., 2017). As such, we observed that YAP/TAZ depletion significantly decreased the number and size of vinculin puncta and nuclear size (**Fig. 5L-N; Suppl. 8G**).

Collectively, these data demonstrate that YAP/TAZ depletion using siRNA leads to impaired YAP/TAZ signaling, FA formation, cytoskeletal/nuclear and ECM remodeling, and cell contractile properties.

### 3.6 YAP/TAZ-TEAD interaction is required for YAP/TAZ downstream effects

Canonically, YAP/TAZ bind to TEAD family members to induce the transcription of YAP/TAZ target genes (Low et al., 2014). To test the effects of YAP/TAZ-TEAD interaction on HTM cell behavior under simulated glaucomatous conditions, we tested the effects of verteporfin (VP), a selective inhibitor of the YAP/TAZ-TEAD transcriptional complex (Liu-Chittenden et al., 2012). HTM cells were seeded on top of or encapsulated in soft hydrogels, and treated with TGFβ2 ± VP. Interestingly, we observed that HTM cells inside hydrogels showed qualitatively decreased YAP nuclear-to-cytoplasmic ratio compared to cells atop of the hydrogels in absence of any treatments (**Fig. 6A-D**). In both cell culture environments, TGFβ2 increased nuclear YAP/TAZ localization, and treating HTM cells with VP significantly decreased TGFβ2-induced nuclear YAP/TAZ as well as downstream TGM2 compared to controls, consistent with the effects of siRNA-mediated YAP/TAZ depletion (**Fig. 6A-D; Suppl. Fig. 9A-D**). We observed that the inhibition of YAP/TAZ-TEAD interaction not only decreased nuclear YAP/TAZ, but also reduced cytoplasmic and total protein levels (**Suppl. Fig. 9E**). VP treatment significantly decreased TGFβ2-induced F-actin filaments, expression of αSMA and p-MLC, and FN deposition (**Fig. 6E-G; Suppl. Fig. 9F-K**).

**Fig. 6.**
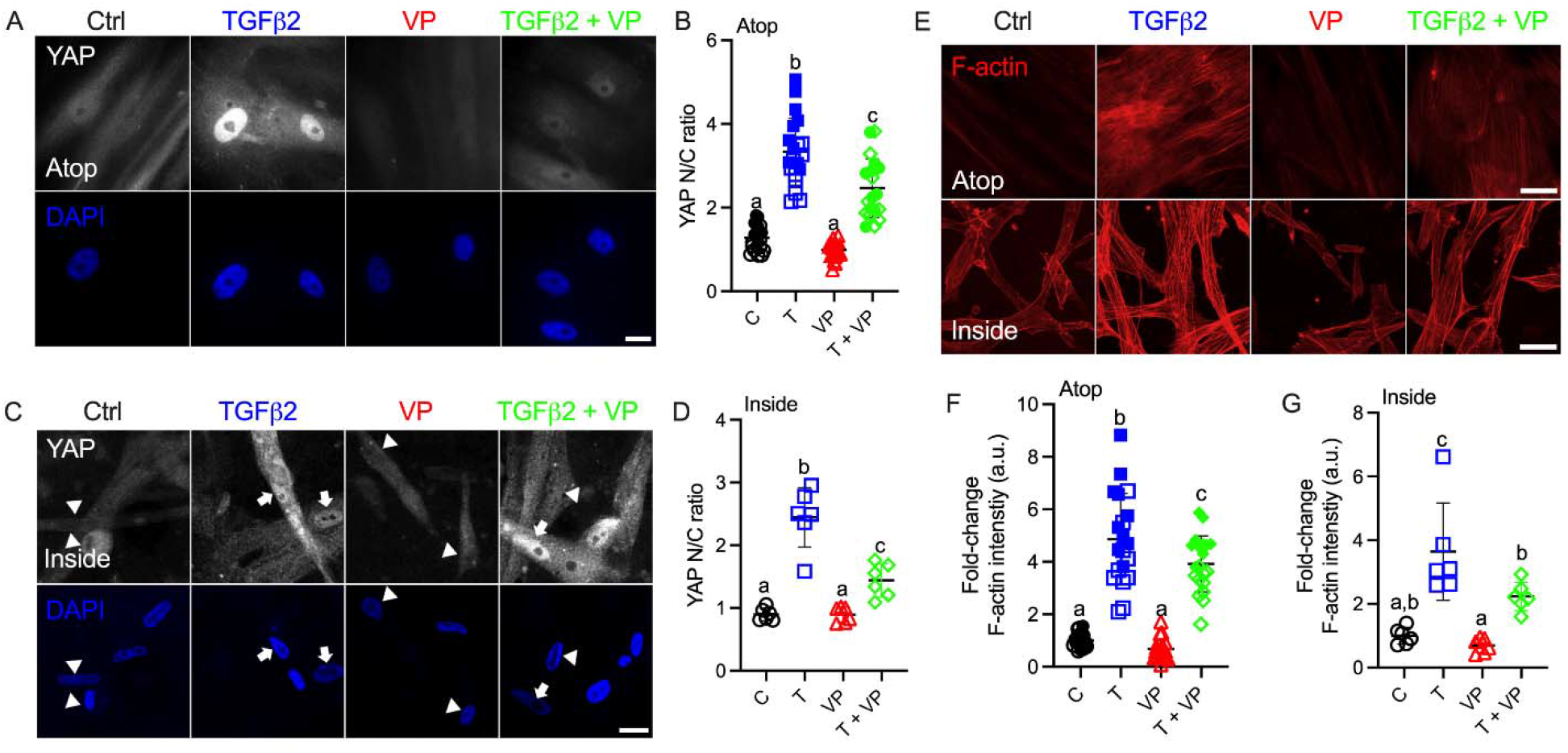
Inhibition of YAP/TAZ-TEAD interaction with VP decreases nuclear YAP and F-actin levels. (A) Representative fluorescence micrographs of YAP in HTM cells cultured on top of soft hydrogels subjected to control, TGFβ2 (2.5 ng/mL), VP (0.5 μM), TGFβ2 + VP at 3 d (YAP = grey; DAPI = blue). Scale bar, 20 μm. (B) Analysis of YAP nuclear/cytoplasmic ratio (N = 20 images from 2 HTM cell strains with 3 replicates per cell strain). (C) Representative fluorescence micrographs of YAP in HTM cells encapsulated in soft hydrogels subjected to the different treatments at 3d (YAP = grey; DAPI = blue). Arrows indicate higher YAP in nuclei, arrowheads indicate higher YAP in cytoplasm. Scale bar, 20 μm. (D) Analysis of YAP nuclear/cytoplasmic ratio (N = 6 images from one HTM cell strain with 3 replicates). (E) Representative fluorescence micrographs of F-actin in HTM cells on or inside of soft hydrogels subjected to the different treatments (F-actin = red). Scale bar, 50 μm. (F) Analysis of F-actin intensity in HTM cells on soft hydrogels (N = 20 images from 2 HTM cell strains with 3 replicates per cell strain). (G) Analysis of F-actin intensity in HTM cells inside of soft hydrogels (N = 6 images from one HTM cell strain with 3 replicates). Open and closed symbols represent different cell strains. The bars and error bars indicate Mean ± SD; Significance was determined by one-way ANOVA using multiple comparisons tests (shared significance indicator letters represent non-significant difference (p>0.05), distinct letters represent significant difference (p<0.05)).

Collectively, these results show that pharmacological inhibition of YAP/TAZ-TEAD interaction leads to impaired ECM remodeling and cell contractile properties, which is independent of cell culture environments (atop or encapsulated in ECM hydrogels).

### 3.7 YAP and TAZ mediate HTM cell contractility and HTM hydrogel stiffness

Lastly, to investigate the effects of YAP/TAZ on HTM cell contractility and ECM stiffening, we encapsulated HTM cells in ECM hydrogels and treated with TGFβ2, either alone or in combination with VP, and assessed hydrogel contractility and stiffness. TGFβ2-treated HTM hydrogels exhibited significantly greater contraction vs. controls by 5 d (74.30% of controls), consistent with our previous report (Li et al., 2021a). Co-treatment of TGFβ2 + VP potently decreased HTM hydrogel contraction (90.37% of controls) compared to TGFβ2-treated samples (**Fig. 7A**). We observed that TGFβ2 significantly increased hydrogel stiffness (1.86-fold of controls), which was prevented by co-treatment with VP (1.16-fold of controls) (**Fig. 7B**).

**Fig. 7.**
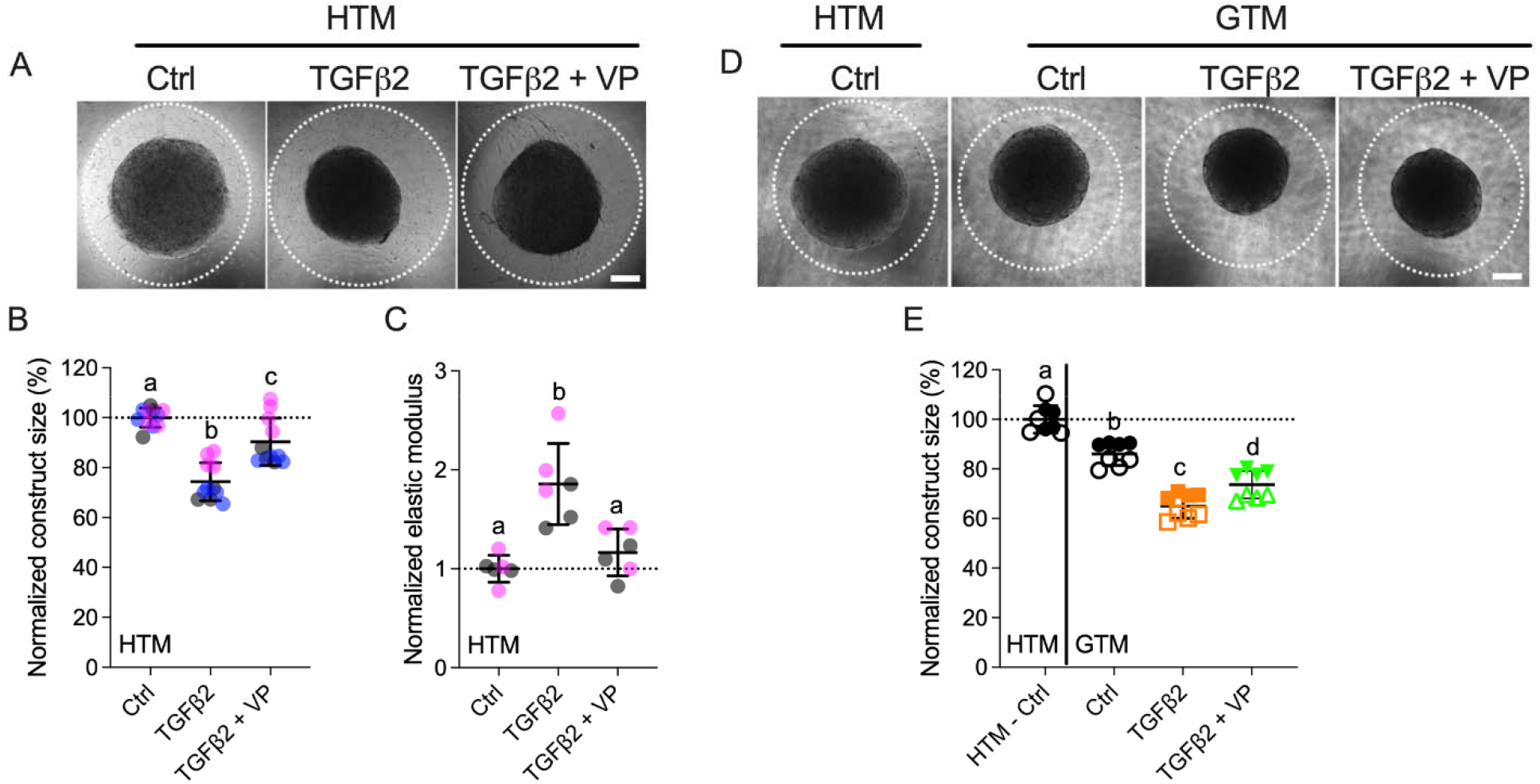
Inhibition of YAP/TAZ activity decreases HTM hydrogel contractility and stiffness. (A) Representative brightfield images of HTM hydrogels subjected to control, TGFβ2 (2.5 ng/ml), TGFβ2 + VP (0.5 μM) at 5 d (dashed lines outline original size of constructs at 0 d). Scale bar, 1 mm. (B) Construct size quantification of HTM hydrogels subjected to the different treatments (N = 11 replicates per group from 3 HTM cell strains). (C) Normalized elastic modulus (to controls) of HTM hydrogels subjected to the different treatments (N = 6 replicates per group from 3 HTM cell strains). (D) Representative brightfield images of GTM hydrogels subjected to the different treatments at 5 d (dashed lines outline original size of constructs at 0 d). Scale bar, 1 mm. (E) Construct size quantification of GTM hydrogels subjected to the different treatments (N = 8 replicates per group from 2 GTM cell strains). Open and closed symbols, or symbols with different colors represent different cell strains; dotted line shows control value for reference. The bars and error bars indicate Mean ± SD; Significance was determined by one-way ANOVA using multiple comparisons tests (shared significance indicator letters represent non-significant difference (p>0.05), distinct letters represent significant difference (p<0.05)).

To assess whether YAP/TAZ inhibition had comparable effects on GTM cells, we evaluated GTM cell-laden hydrogel contraction in response to the same treatments. Consistent with our previous study (Li et al., 2021a), we demonstrated that GTM hydrogels in absence of additional TGFβ2 induction exhibited significantly greater contraction relative to normal HTM hydrogels (86.08% of HTM hydrogels controls); TGFβ2 further increased GTM hydrogel contraction, and VP partially blocked this during the short 5 d exposure (**Fig. 7C**).

To determine if hydrogel contractility was influenced by the cell number, we assessed HTM/GTM cell proliferation in constructs subjected to the different treatments. We observed a fewer number of cells in the TGFβ2 + VP group compared to TGFβ2-treated samples (10.10% decreased) (**Suppl. Fig. 10A**), while co-treatment of TGFβ2 + VP reduced hydrogel contraction and stiffening by 21.63% and 59.45% compared to the TGFβ2-treated group, respectively (**Fig. 7A,B**); demonstrating that VP induced decreasing hydrogel contraction was not only caused by the smaller cell number, but also reduced cell contractility. No significant differences between the different groups were observed for GTM cell-laden hydrogels (**Suppl. Fig. 10B**).

Together, these data demonstrate that TGFβ2 robustly induces HTM hydrogel contractility and stiffening in a soft ECM environment, which are potently reduced by YAP/TAZ inhibition. Likewise, inhibition of YAP/TAZ had similar effects on GTM cells inside the 3D hydrogel network.

## 4. Discussion

The mechanosensitive transcriptional coactivators YAP and TAZ play important roles in mechanotransduction, a process through which cells translate external biophysical cues into internal biochemical signals. YAP/TAZ modulate target gene expression profiles with broad functional consequences across many cell and tissue types (Dupont et al., 2011; Boopathy and Hong, 2019). Through this mechanism, YAP/TAZ signaling regulates critical cellular functions and normal tissue homeostasis; imbalance or failure of this process is at the core of various diseases (Panciera et al., 2017). Indeed, elevated YAP/TAZ transcriptional activity is associated with glaucomatous HTM cell dysfunction (Thomasy et al., 2013; Chen et al., 2015; Ho et al., 2018; Peng et al., 2018; Dhamodaran et al., 2020; Yemanyi and Raghunathan, 2020; Yemanyi et al., 2020a). Importantly, a recent genome-wide meta-analysis identified YAP among 44 previously unknown POAG risk loci (Gharahkhani et al., 2021a); this observation provides strong new evidence that YAP may play a prominent role in glaucoma pathogenesis. Here, we found that YAP/TAZ activity is markedly upregulated in TM cells from patients with glaucoma compared to cells isolated from healthy tissue, exhibiting normal variability between cells from different donors (i.e., one GTM cell strain showed higher baseline YAP/TAZ nuclear localization and transcriptional activity compared to the other) (**Fig. 1; Suppl. Fig. 2,3**). However, the detailed mechanisms for YAP/TAZ modulation in HTM cells under glaucomatous conditions (i.e., stiffened ECM and increased growth factors in AH) remain to be elucidated. To model this, we used biomimetic ECM hydrogels with tunable stiffness to study the roles of YAP and TAZ in HTM cells in response to stiffened matrix and TGFβ2. As summarized in **Fig. 8**, our data support that YAP/TAZ are critical regulators in mediating HTM cellular responses to the stiffened ECM and elevated TGFβ2 in POAG; we propose that increased YAP/TAZ activity may drive further HTM tissue stiffening to exacerbate disease pathology conditions. This conclusion is supported by the findings that (i) stiffened ECM hydrogels elevate YAP/TAZ activity, as indicated by nuclear localization of YAP/TAZ, potentially through regulating FA formation and cytoskeleton rearrangement; (ii) TGFβ2 induces nuclear YAP/TAZ localization and target gene activation through ERK and ROCK signaling pathways; (iii) depolymerization of F-actin decreases YAP/TAZ activity; (iv) YAP/TAZ depletion using siRNA or pharmacological inhibition of YAP/TAZ-TEAD interaction decreases FA formation, cytoskeletal/nuclear/ECM remodeling and cell contractile properties; (v) YAP/TAZ inhibitor VP decreases HTM/GTM cell-laden hydrogel contraction and HTM hydrogel stiffening.

**Fig. 8.**
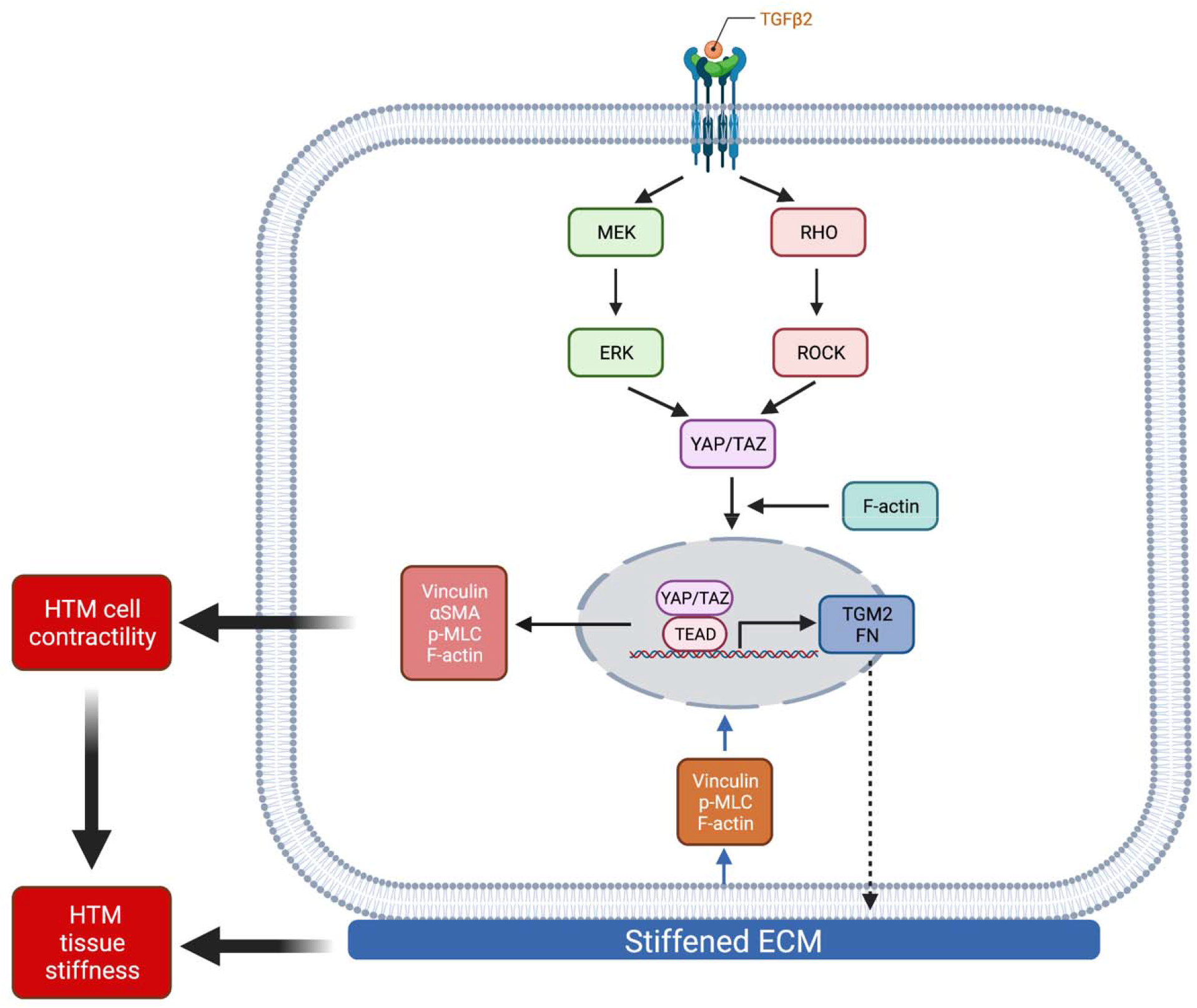
Schematic illustration of the effects elicited by ECM stiffness and TGFβ2 that modulate YAP/TAZ activity in HTM cells. Stiffened ECM hydrogels elevate YAP/TAZ activity potentially through regulating focal adhesion formation and actin cytoskeleton rearrangement. TGFβ2 activates ERK and ROCK signaling pathways to regulate YAP/TAZ activity in normal HTM and GTM cells. YAP/TAZ activation induces HTM cell contractility and ECM remodeling, which together may increase HTM stiffness in POAG. Created with BioRender.com.

Most *in vitro* studies of HTM cell (patho-)physiology have relied on conventional cell monolayer cultures on plastic or glass of supraphysiologic stiffness. However, biophysical cues such as substrate composition and stiffness are known to be potent modulators of cell behaviors (Raghunathan et al., 2013). In our previous studies, we reported that HTM cells on glass exhibited bigger nuclei and different organization of F-actin fibers compared to HTM cells on ECM hydrogels (Li et al., 2021b). Here, we demonstrated that HTM cells on hydrogels showed a smaller number of vinculin FA vs. HTM cells on glass (**Fig. 2C; Suppl. Fig. 5A**), confirming that HTM cells display distinct physiological characteristics on soft tissue-mimetic biomaterials and traditional tissue culture plastic or glass substrates of supraphysiological stiffness (Caliari and Burdick, 2016).

Increased tissue stiffness has been observed in multiple pathologies, including cancer, cardiovascular and fibrosis-related diseases (Lampi and Reinhart-King, 2018). ECM stiffening can precede disease development and consequently increased mechanical cues can drive their progression via altered mechanotransduction (Frantz et al., 2010; Kaess et al., 2012; Pickup et al., 2014). Therefore, therapeutically targeting ECM stiffening by disrupting the cellular response to the stiffened ECM environment, or in other words targeting mechanotransduction, is an emerging field with clear implications for glaucoma treatment. It is widely accepted that the ECM is stiffer in the glaucomatous HTM (Wang et al., 2017b). Previous studies have also shown that ECM deposited by HTM cells treated with DEX was ∼2-4-fold stiffer relative to controls (Raghunathan et al., 2015; Raghunathan et al., 2018). Our recent data using DEX-induced HTM cell-encapsulated 3D ECM hydrogels were in good agreement with these observations (Li et al., 2021a). In this study, we demonstrated that TGFβ2 treatment resulted in a comparable ∼2-fold increase in ECM stiffness surrounding the HTM cells as they reside embedded in our bioengineered hydrogels (**Fig. 2A**). In both scenarios, the stiffness changes were cell-driven in response to biochemical cues implicated in glaucoma.

Corneal UV crosslinking with riboflavin (RF) is a clinical treatment to stabilize the collagen-rich stroma in corneal ectasias (Zhang et al., 2011; Rahman et al., 2020), with promising potential for enhancing mechanical properties of collagen-based hydrogels (Ahearne and Coyle, 2016; Heo et al., 2016). Crosslinking occurs via covalent bond formation between amino acids of collagen fibrils induced by singlet O_2_ from UV-excited RF (Tirella et al., 2012). To simulate glaucomatous ECM stiffening for investigations of the cellular response, we used riboflavin to double-crosslink collagen in the hydrogels, which increased their stiffness ∼2-fold over baseline (**Fig. 2B**). HTM cells on the stiffened hydrogels exhibited bigger nuclei, increased number of vinculin FA, cytoskeleton rearrangement, ECM deposition and YAP/TAZ nuclear localization compared to cells on soft hydrogels (**Fig. 2C-L; Suppl. Fig.3**). Taken together, our findings indicate that YAP/TAZ activity in HTM cells is distinctive on different substrates, suggesting that hydrogels with tunable physiochemical properties may be more suitable to investigate subtleties in HTM cell behaviors that would otherwise go unnoticed when relying exclusively on traditional stiff 2D culture substrates.

TGFβ2 levels are elevated in the aqueous humor of glaucoma patients compared to age-matched normal eyes (Inatani et al., 2001; Picht et al., 2001; Ochiai and Ochiai, 2002; Agarwal et al., 2015). Here, we showed that GTM cells isolated from POAG donor eyes secreted significantly more active TGFβ2 compared to normal HTM cells (**Fig. 1A; Suppl. Fig. 2**). We observed that TGFβ2 upregulated YAP/TAZ nuclear localization and TGM2 expression, a downstream effector of active YAP/TAZ signaling, in both normal HTM and GTM cells, confirming that YAP/TAZ activity are upregulated under glaucomatous conditions. ERK or ROCK inhibition decreased nuclear YAP/TAZ and TGM2 in both HTM and GTM cells, and co-treatment of ERK or ROCK inhibitor with TGFβ2 restored YAP/TAZ cellular localization and TGM2 to HTM control levels (**Fig. 3; Suppl. Fig. 6**). Further research will be necessary to investigate in greater detail whether/how ERK and ROCK signaling differentially regulate YAP and TAZ activity in HTM cells.

Actomyosin cell contractility forces are increased in response to elevated ECM stiffness and TGFβ2 induction (Lampi and Reinhart-King, 2018; Li et al., 2021b). Additionally, we have demonstrated that the stiffened ECM induces YAP/TAZ nuclear localization, and increases F-actin filaments, p-MLC and αSMA - all involved in actomyosin cell contractility force generation. We hypothesized that increased actomyosin cell contractility may drive YAP/TAZ nuclear localization in HTM cells. Here, we used Lat B to depolymerize F-actin stress fibers and increase cytoskeletal relaxation. We found that a short exposure time to Lat B eliminated vinculin FA formation and YAP/TAZ nuclear localization, and decreased TGM2 expression. ROCK inhibition has also been shown to decrease HTM cell F-actin fibers and contractility (Li et al., 2021a). These observations on effects of Lat B were consistent with our findings that ROCK inhibitor reduced YAP/TAZ activity (**Fig. 3; Fig. 4F-J; Suppl. Fig. 7**). It would be worthwhile to further investigate effects of other cell contractility related molecules, such as myosin light chain kinase, myosin II, cofilin and gelsolin, on YAP/TAZ activity in both HTM and GTM cells.

We have observed that simulated glaucomatous conditions (i.e., elevated TGFβ2 and stiffened ECM) upregulated YAP/TAZ activity, and treatments that can reduce IOP in experimental models (i.e., ROCK inhibitor and Lat B) downregulated YAP/TAZ activity. These observations led us to explore the role of YAP/TAZ on glaucoma pathology development. We found that YAP/TAZ depletion using siRNA consistently decreased FA formation (i.e., vinculin), and reduced expression of cell contractile proteins (i.e., F-actin, p-MLC and αSMA) and ECM proteins (i.e., FN) (**Fig. 5**). Also, YAP/TAZ deactivation reduced expression of TGM2, a protein that promotes cell-matrix interactions and FN crosslinking to stiffen the ECM (Akimov et al., 2000), and CTGF, known to increase ECM production and cell contractility in HTM cells, and elevate IOP in mouse eyes (Junglas et al., 2012). Thus, YAP/TAZ inhibition may decrease subsequent ECM stiffness potentially through regulation of ECM, TGM2 and CTGF production. Furthermore, we demonstrated that inhibition of YAP/TAZ-TEAD interaction using VP significantly decreased TGFβ2-induced YAP/TAZ nuclear localization in HTM cells independent of their spatial arrangement atop or inside of the ECM hydrogels. Consistent with the effects of siRNA-mediated YAP/TAZ depletion, VP treatment rescued TGFβ2-induced YAP/TAZ activity, cell contractile properties and ECM remodeling (**Fig. 6; Suppl. Fig. 9**).

Notably, YAP/TAZ inhibition using VP blunted HTM/GTM cell-laden hydrogel contraction and stiffening (**Fig. 7**). Thus, we conclude that YAP/TAZ act as central players in regulating HTM cell mechanical homeostasis in response to changes of the surrounding microenvironment (e.g., levels of growth factors, ECM stiffness) to maintain tissue-level structural integrity and functionality. It has been demonstrated that the relative roles of YAP/TAZ are cell type- and context-dependent; they can cause homeostatic regulation of tissue properties (negative feedback loop) or promote fibrotic conditions (positive feedback loop). Some reports implicated YAP/TAZ in a feed-forward promotion of cytoskeletal tension and ECM protein deposition (Lin et al., 2017; Nardone et al., 2017; Yemanyi and Raghunathan, 2020), while some research showed YAP/TAZ had a negative feedback regulation that acted to suppress actin polymerization and cytoskeletal tension (Qiao et al., 2017; Mason et al., 2019). Our data were consistent with the former; i.e., YAP/TAZ drives HTM cell contraction and ECM stiffening.

In conclusion, using our bioengineered tissue-mimetic ECM hydrogel system, we demonstrated that YAP/TAZ activity is upregulated in response to simulated glaucomatous conditions (i.e., TGFβ2 induction and stiffened ECM), and that YAP/TAZ activation induces HTM cell contractility and ECM remodeling, which together may increase HTM stiffness in POAG. Our findings provide strong evidence for a pathologic role of aberrant YAP/TAZ signaling in glaucomatous HTM cell dysfunction, and may help inform strategies for the development of novel multifactorial approaches to prevent progressive ocular hypertension in glaucoma.

## Acknowledgments

We thank Dr. Robert W. Weisenthal and the team at Specialty Surgery Center of Central New York for assistance with corneal rim specimens. We also thank Dr. Nasim Annabi at the University of California – Los Angeles for providing the KCTS-ELP, Dr. Alison Patteson at Syracuse University for rheometer access, Drs. Audrey M. Bernstein and Mariano S. Viapiano, and Neuroscience Microscopy Core at Upstate Medical University for imaging support.

## Author contributions

H.L., VK.R., W.D.S., P.S.G., and S.H. designed all experiments, collected, analyzed, and interpreted the data. W.D.S. provided the GTM cells. VK.R. performed the AFM experiments. H.L. and S.H. wrote the manuscript. All authors commented on and approved the final manuscript. P.S.G. and S.H. conceived and supervised the research.

## Data and materials availability

All data needed to evaluate the conclusions in the paper are present in the paper and/or the Supplementary Materials. Additional data related to this paper may be requested from the authors.

## References

Abu-Hassan, D.W., Acott, T.S., and Kelley, M.J. (2014). The Trabecular Meshwork: A Basic Review of Form and Function. J Ocul Biol 2(1), 9.

Acott, T.S., and Kelley, M.J. (2008). Extracellular matrix in the trabecular meshwork. Exp Eye Res 86(4), 543–561. doi: 10.1016/j.exer.2008.01.013.

Agarwal, P., Daher, A.M., and Agarwal, R. (2015). Aqueous humor TGF-β2 levels in patients with open-angle glaucoma: A meta-analysis. Molecular vision 21, 612–620.

Ahearne, M., and Coyle, A. (2016). Application of UVA-riboflavin crosslinking to enhance the mechanical properties of extracellular matrix derived hydrogels. J Mech Behav Biomed Mater 54, 259–267. doi: 10.1016/j.jmbbm.2015.09.035.

Akimov, S.S., Krylov, D., Fleischman, L.F., and Belkin, A.M. (2000). Tissue transglutaminase is an integrin-binding adhesion coreceptor for fibronectin. The Journal of cell biology 148(4), 825–838. doi: 10.1083/jcb.148.4.825.

Boopathy, G.T.K., and Hong, W. (2019). Role of Hippo Pathway-YAP/TAZ Signaling in Angiogenesis. Frontiers in Cell and Developmental Biology 7(49). doi: 10.3389/fcell.2019.00049.

Brubaker, R.F. (1991). Flow of aqueous humor in humans [The Friedenwald Lecture]. Invest Ophthalmol Vis Sci 32(13), 3145–3166.

Caliari, S.R., and Burdick, J.A. (2016). A practical guide to hydrogels for cell culture. Nat Methods 13(5), 405–414. doi: 10.1038/nmeth.3839.

Chang, Y.R., Raghunathan, V.K., Garland, S.P., Morgan, J.T., Russell, P., and Murphy, C.J. (2014). Automated AFM force curve analysis for determining elastic modulus of biomaterials and biological samples. J Mech Behav Biomed Mater 37, 209–218. doi: 10.1016/j.jmbbm.2014.05.027.

Chen, W.S., Cao, Z., Krishnan, C., and Panjwani, N. (2015). Verteporfin without light stimulation inhibits YAP activation in trabecular meshwork cells: Implications for glaucoma treatment. Biochem Biophys Res Commun 466(2), 221–225. doi: 10.1016/j.bbrc.2015.09.012.

Dhamodaran, K., Baidouri, H., Sandoval, L., and Raghunathan, V. (2020). Wnt Activation After Inhibition Restores Trabecular Meshwork Cells Toward a Normal Phenotype. Invest Ophthalmol Vis Sci 61(6), 30. doi: 10.1167/iovs.61.6.30.

Drury, J.L., and Mooney, D.J. (2003). Hydrogels for tissue engineering: scaffold design variables and applications. Biomaterials 24(24), 4337–4351. doi: https://doi.org/10.1016/S0142-9612(03)00340-5.

Dupont, S., Morsut, L., Aragona, M., Enzo, E., Giulitti, S., Cordenonsi, M., et al. (2011). Role of YAP/TAZ in mechanotransduction. Nature 474(7350), 179–183. doi: 10.1038/nature10137.

Ethier, C.R., Read, A.T., and Chan, D.W. (2006). Effects of latrunculin-B on outflow facility and trabecular meshwork structure in human eyes. Invest Ophthalmol Vis Sci 47(5), 1991–1998. doi: 10.1167/iovs.05-0327.

Frantz, C., Stewart, K.M., and Weaver, V.M. (2010). The extracellular matrix at a glance. J Cell Sci 123(Pt 24), 4195–4200. doi: 10.1242/jcs.023820.

Fuchshofer, R., and Tamm, E.R. (2009). Modulation of extracellular matrix turnover in the trabecular meshwork. Experimental Eye Research 88(4), 683–688. doi: https://doi.org/10.1016/j.exer.2009.01.005.

Gharahkhani, P., Jorgenson, E., Hysi, P., Khawaja, A.P., Pendergrass, S., Han, X., et al. (2021a). Genome-wide meta-analysis identifies 127 open-angle glaucoma loci with consistent effect across ancestries. Nat Commun 12(1), 1258. doi: 10.1038/s41467-020-20851-4.

Gharahkhani, P., Jorgenson, E., Hysi, P., Khawaja, A.P., Pendergrass, S., Han, X., et al. (2021b). Genome-wide meta-analysis identifies 127 open-angle glaucoma loci with consistent effect across ancestries. Nat Commun 12(1), 1258. doi: 10.1038/s41467-020-20851-4.

Granstein, R.D., Staszewski, R., Knisely, T.L., Zeira, E., Nazareno, R., Latina, M., et al. (1990). Aqueous humor contains transforming growth factor-beta and a small (less than 3500 daltons) inhibitor of thymocyte proliferation. J Immunol 144(8), 3021–3027.

Han, H., Wecker, T., Grehn, F., and Schlunck, G. (2011). Elasticity-Dependent Modulation of TGF-β Responses in Human Trabecular Meshwork Cells. Investigative Ophthalmology & Visual Science 52(6), 2889–2896. doi: 10.1167/iovs.10-6640.

Hann, C.R., and Fautsch, M.P. (2011). The elastin fiber system between and adjacent to collector channels in the human juxtacanalicular tissue. Invest Ophthalmol Vis Sci 52(1), 45–50. doi: 10.1167/iovs.10-5620.

Heo, J., Koh, R.H., Shim, W., Kim, H.D., Yim, H.G., and Hwang, N.S. (2016). Riboflavin-induced photo-crosslinking of collagen hydrogel and its application in meniscus tissue engineering. Drug Deliv Transl Res 6(2), 148–158. doi: 10.1007/s13346-015-0224-4.

Ho, L.T.Y., Skiba, N., Ullmer, C., and Rao, P.V. (2018). Lysophosphatidic Acid Induces ECM Production via Activation of the Mechanosensitive YAP/TAZ Transcriptional Pathway in Trabecular Meshwork Cells. Invest Ophthalmol Vis Sci 59(5), 1969–1984. doi: 10.1167/iovs.17-23702.

Honjo, M., Igarashi, N., Nishida, J., Kurano, M., Yatomi, Y., Igarashi, K., et al. (2018). Role of the Autotaxin-LPA Pathway in Dexamethasone-Induced Fibrotic Responses and Extracellular Matrix Production in Human Trabecular Meshwork Cells. Investigative Ophthalmology & Visual Science 59(1), 21–30. doi: 10.1167/iovs.17-22807.

Inatani, M., Tanihara, H., Katsuta, H., Honjo, M., Kido, N., and Honda, Y. (2001). Transforming growth factor-beta 2 levels in aqueous humor of glaucomatous eyes. Graefes Arch Clin Exp Ophthalmol 239(2), 109–113. doi: 10.1007/s004170000241.

Jansen, K.A., Atherton, P., and Ballestrem, C. (2017). Mechanotransduction at the cell-matrix interface. Semin Cell Dev Biol 71, 75–83. doi: 10.1016/j.semcdb.2017.07.027.

Junglas, B., Kuespert, S., Seleem, A.A., Struller, T., Ullmann, S., Bösl, M., et al. (2012). Connective Tissue Growth Factor Causes Glaucoma by Modifying the Actin Cytoskeleton of the Trabecular Meshwork. The American Journal of Pathology 180(6), 2386–2403. doi: https://doi.org/10.1016/j.ajpath.2012.02.030.

Kaess, B.M., Rong, J., Larson, M.G., Hamburg, N.M., Vita, J.A., Levy, D., et al. (2012). Aortic stiffness, blood pressure progression, and incident hypertension. Jama 308(9), 875–881. doi: 10.1001/2012.jama.10503.

Kasetti, R.B., Maddineni, P., Patel, P.D., Searby, C., Sheffield, V.C., and Zode, G.S. (2018). Transforming growth factor β2 (TGFβ2) signaling plays a key role in glucocorticoid-induced ocular hypertension. J Biol Chem 293(25), 9854–9868. doi: 10.1074/jbc.RA118.002540.

Keller, K.E., and Acott, T.S. (2013). The Juxtacanalicular Region of Ocular Trabecular Meshwork: A Tissue with a Unique Extracellular Matrix and Specialized Function. J Ocul Biol 1(1), 3.

Keller, K.E., Bhattacharya, S.K., Borras, T., Brunner, T.M., Chansangpetch, S., Clark, A.F., et al. (2018). Consensus recommendations for trabecular meshwork cell isolation, characterization and culture. Exp Eye Res 171, 164–173. doi: 10.1016/j.exer.2018.03.001.

Kwon, Y.H., Fingert, J.H., Kuehn, M.H., and Alward, W.L. (2009). Primary open-angle glaucoma. N Engl J Med 360(11), 1113–1124. doi: 10.1056/NEJMra0804630.

Lampi, M.C., and Reinhart-King, C.A. (2018). Targeting extracellular matrix stiffness to attenuate disease: From molecular mechanisms to clinical trials. Sci Transl Med 10(422). doi: 10.1126/scitranslmed.aao0475.

Last, J.A., Pan, T., Ding, Y., Reilly, C.M., Keller, K., Acott, T.S., et al. (2011). Elastic Modulus Determination of Normal and Glaucomatous Human Trabecular Meshwork. Investigative Ophthalmology & Visual Science 52(5), 2147–2152. doi: 10.1167/iovs.10-6342.

Lei, Q.Y., Zhang, H., Zhao, B., Zha, Z.Y., Bai, F., Pei, X.H., et al. (2008). TAZ promotes cell proliferation and epithelial-mesenchymal transition and is inhibited by the hippo pathway. Mol Cell Biol 28(7), 2426–2436. doi: 10.1128/mcb.01874-07.

Li, H., Bague, T., Kirschner, A., Strat, A.N., Roberts, H., Weisenthal, R.W., et al. (2021a). A tissue-engineered human trabecular meshwork hydrogel for advanced glaucoma disease modeling. Exp Eye Res 205, 108472. doi: 10.1016/j.exer.2021.108472.

Li, H., Henty-Ridilla, J.L., Ganapathy, P.S., and Herberg, S. (2021b). TGFβ2 regulates human trabecular meshwork cell contractility via ERK and ROCK pathways with distinct signaling crosstalk dependent on the culture substrate. bioRxiv, 2021.2007.2001.450718. doi: 10.1101/2021.07.01.450718.

Lin, C., Yao, E., Zhang, K., Jiang, X., Croll, S., Thompson-Peer, K., et al. (2017). YAP is essential for mechanical force production and epithelial cell proliferation during lung branching morphogenesis. Elife 6. doi: 10.7554/eLife.21130.

Liu-Chittenden, Y., Huang, B., Shim, J.S., Chen, Q., Lee, S.J., Anders, R.A., et al. (2012). Genetic and pharmacological disruption of the TEAD-YAP complex suppresses the oncogenic activity of YAP. Genes Dev 26(12), 1300–1305. doi: 10.1101/gad.192856.112.

Low, B.C., Pan, C.Q., Shivashankar, G.V., Bershadsky, A., Sudol, M., and Sheetz, M. (2014). YAP/TAZ as mechanosensors and mechanotransducers in regulating organ size and tumor growth. FEBS Lett 588(16), 2663–2670. doi: 10.1016/j.febslet.2014.04.012.

Ma, J., Sanchez-Duffhues, G., Goumans, M.-J., and Ten Dijke, P. (2020). TGF-β-Induced Endothelial to Mesenchymal Transition in Disease and Tissue Engineering. Frontiers in cell and developmental biology 8, 260–260. doi: 10.3389/fcell.2020.00260.

Mason, D.E., Collins, J.M., Dawahare, J.H., Nguyen, T.D., Lin, Y., Voytik-Harbin, S.L., et al. (2019). YAP and TAZ limit cytoskeletal and focal adhesion maturation to enable persistent cell motility. J Cell Biol 218(4), 1369–1389. doi: 10.1083/jcb.201806065.

McKee, C.T., Wood, J.A., Shah, N.M., Fischer, M.E., Reilly, C.M., Murphy, C.J., et al. (2011). The effect of biophysical attributes of the ocular trabecular meshwork associated with glaucoma on the cell response to therapeutic agents. Biomaterials 32(9), 2417–2423. doi: https://doi.org/10.1016/j.biomaterials.2010.11.071.

Mohammed, D., Versaevel, M., Bruyère, C., Alaimo, L., Luciano, M., Vercruysse, E., et al. (2019). Innovative Tools for Mechanobiology: Unraveling Outside-In and Inside-Out Mechanotransduction. Frontiers in Bioengineering and Biotechnology 7(162). doi: 10.3389/fbioe.2019.00162.

Montecchi-Palmer, M., Bermudez, J.Y., Webber, H.C., Patel, G.C., Clark, A.F., and Mao, W. (2017). TGFβ2 Induces the Formation of Cross-Linked Actin Networks (CLANs) in Human Trabecular Meshwork Cells Through the Smad and Non-Smad Dependent Pathways. Investigative ophthalmology & visual science 58(2), 1288–1295. doi: 10.1167/iovs.16-19672.

Muppala, S., Raghunathan, V.K., Jalilian, I., Thomasy, S., and Murphy, C.J. (2019). YAP and TAZ are distinct effectors of corneal myofibroblast transformation. Exp Eye Res 180, 102–109. doi: 10.1016/j.exer.2018.12.009.

Nardone, G., Oliver-De La Cruz, J., Vrbsky, J., Martini, C., Pribyl, J., Skládal, P., et al. (2017). YAP regulates cell mechanics by controlling focal adhesion assembly. Nature Communications 8(1), 15321. doi: 10.1038/ncomms15321.

Ochiai, Y., and Ochiai, H. (2002). Higher concentration of transforming growth factor-beta in aqueous humor of glaucomatous eyes and diabetic eyes. Jpn J Ophthalmol 46(3), 249–253. doi: 10.1016/s0021-5155(01)00523-8.

Panahi, R., and Baghban-Salehi, M. (2019). “Protein-Based Hydrogels,” in Cellulose-Based Superabsorbent Hydrogels, ed. M.I.H. Mondal. (Cham: Springer International Publishing), 1561–1600.

Panciera, T., Azzolin, L., Cordenonsi, M., and Piccolo, S. (2017). Mechanobiology of YAP and TAZ in physiology and disease. Nat Rev Mol Cell Biol 18(12), 758–770. doi: 10.1038/nrm.2017.87.

Peng, J., Wang, H., Wang, X., Sun, M., Deng, S., and Wang, Y. (2018). YAP and TAZ mediate steroid-induced alterations in the trabecular meshwork cytoskeleton in human trabecular meshwork cells. Int J Mol Med 41(1), 164–172. doi: 10.3892/ijmm.2017.3207.

Peterson, J.A., Tian, B., Bershadsky, A.D., Volberg, T., Gangnon, R.E., Spector, I., et al. (1999). Latrunculin-A increases outflow facility in the monkey. Invest Ophthalmol Vis Sci 40(5), 931–941.

Picht, G., Welge-Luessen, U., Grehn, F., and Lutjen-Drecoll, E. (2001). Transforming growth factor beta 2 levels in the aqueous humor in different types of glaucoma and the relation to filtering bleb development. Graefes Arch Clin Exp Ophthalmol 239(3), 199–207. doi: 10.1007/s004170000252.

Pickup, M.W., Mouw, J.K., and Weaver, V.M. (2014). The extracellular matrix modulates the hallmarks of cancer. EMBO Rep 15(12), 1243–1253. doi: 10.15252/embr.201439246.

Prendes, M.A., Harris, A., Wirostko, B.M., Gerber, A.L., and Siesky, B. (2013). The role of transforming growth factor β in glaucoma and the therapeutic implications. Br J Ophthalmol 97(6), 680–686. doi: 10.1136/bjophthalmol-2011-301132.

Qiao, Y., Chen, J., Lim, Y.B., Finch-Edmondson, M.L., Seshachalam, V.P., Qin, L., et al. (2017). YAP Regulates Actin Dynamics through ARHGAP29 and Promotes Metastasis. Cell Rep 19(8), 1495–1502. doi: 10.1016/j.celrep.2017.04.075.

Quigley, H.A. (1993). Open-angle glaucoma. N Engl J Med 328(15), 1097–1106. doi: 10.1056/nejm199304153281507.

Quigley, H.A., and Broman, A.T. (2006). The number of people with glaucoma worldwide in 2010 and 2020. Br J Ophthalmol 90(3), 262–267. doi: 10.1136/bjo.2005.081224.

Raghunathan, V.K., Benoit, J., Kasetti, R., Zode, G., Salemi, M., Phinney, B.S., et al. (2018). Glaucomatous cell derived matrices differentially modulate non-glaucomatous trabecular meshwork cellular behavior. Acta Biomater 71, 444–459. doi: 10.1016/j.actbio.2018.02.037.

Raghunathan, V.K., Morgan, J.T., Dreier, B., Reilly, C.M., Thomasy, S.M., Wood, J.A., et al. (2013). Role of substratum stiffness in modulating genes associated with extracellular matrix and mechanotransducers YAP and TAZ. Invest Ophthalmol Vis Sci 54(1), 378–386. doi: 10.1167/iovs.12-11007.

Raghunathan, V.K., Morgan, J.T., Park, S.A., Weber, D., Phinney, B.S., Murphy, C.J., et al. (2015). Dexamethasone Stiffens Trabecular Meshwork, Trabecular Meshwork Cells, and Matrix. Investigative ophthalmology & visual science 56(8), 4447–4459. doi: 10.1167/iovs.15-16739.

Rahman, N., O’Neill, E., Irnaten, M., Wallace, D., and O’Brien, C. (2020). Corneal Stiffness and Collagen Cross-Linking Proteins in Glaucoma: Potential for Novel Therapeutic Strategy. J Ocul Pharmacol Ther 36(8), 582–594. doi: 10.1089/jop.2019.0118.

Schindelin, J., Arganda-Carreras, I., Frise, E., Kaynig, V., Longair, M., Pietzsch, T., et al. (2012). Fiji: an open-source platform for biological-image analysis. Nat Methods 9(7), 676–682. doi: 10.1038/nmeth.2019.

Schlunck, G.n., Han, H., Wecker, T., Kampik, D., Meyer-ter-Vehn, T., and Grehn, F. (2008). Substrate Rigidity Modulates Cell–Matrix Interactions and Protein Expression in Human Trabecular Meshwork Cells. Investigative Ophthalmology & Visual Science 49(1), 262–269. doi: 10.1167/iovs.07-0956.

Schmittgen, T.D., and Livak, K.J. (2008). Analyzing real-time PCR data by the comparative C(T) method. Nature Protocols 3(6), 1101–1108.

Stamer, W.D., Seftor, R.E., Williams, S.K., Samaha, H.A., and Snyder, R.W. (1995). Isolation and culture of human trabecular meshwork cells by extracellular matrix digestion. Curr Eye Res 14(7), 611–617.

Tamm, E.R. (2009). The trabecular meshwork outflow pathways: structural and functional aspects. Exp Eye Res 88(4), 648–655. doi: 10.1016/j.exer.2009.02.007.

Tamm, E.R., Braunger, B.M., and Fuchshofer, R. (2015). Intraocular Pressure and the Mechanisms Involved in Resistance of the Aqueous Humor Flow in the Trabecular Meshwork Outflow Pathways. Prog Mol Biol Transl Sci 134, 301–314. doi: 10.1016/bs.pmbts.2015.06.007.

Tham, Y.C., Li, X., Wong, T.Y., Quigley, H.A., Aung, T., and Cheng, C.Y. (2014). Global prevalence of glaucoma and projections of glaucoma burden through 2040: a systematic review and meta-analysis. Ophthalmology 121(11), 2081–2090. doi: 10.1016/j.ophtha.2014.05.013.

Thomasy, S.M., Morgan, J.T., Wood, J.A., Murphy, C.J., and Russell, P. (2013). Substratum stiffness and latrunculin B modulate the gene expression of the mechanotransducers YAP and TAZ in human trabecular meshwork cells. Exp Eye Res 113, 66–73. doi: 10.1016/j.exer.2013.05.014.

Tian, B., Geiger, B., Epstein, D.L., and Kaufman, P.L. (2000). Cytoskeletal Involvement in the Regulation of Aqueous Humor Outflow. Investigative Ophthalmology & Visual Science 41(3), 619–623.

Timothy P. Lodge, P.C.H. (2020). Polymer Chemistry. CRC Press. doi: https://doi.org/10.1201/9780429190810.

Tirella, A., Liberto, T., and Ahluwalia, A. (2012). Riboflavin and collagen: New crosslinking methods to tailor the stiffness of hydrogels. Materials Letters 74, 58–61.

Totaro, A., Panciera, T., and Piccolo, S. (2018). YAP/TAZ upstream signals and downstream responses. Nature Cell Biology 20(8), 888–899. doi: 10.1038/s41556-018-0142-z.

Tovar-Vidales, T., Roque, R., Clark, A.F., and Wordinger, R.J. (2008). Tissue transglutaminase expression and activity in normal and glaucomatous human trabecular meshwork cells and tissues. Investigative ophthalmology & visual science 49(2), 622–628. doi: 10.1167/iovs.07-0835.

Vahabikashi, A., Gelman, A., Dong, B., Gong, L., Cha, E.D.K., Schimmel, M., et al. (2019). Increased stiffness and flow resistance of the inner wall of Schlemm’s canal in glaucomatous human eyes. Proceedings of the National Academy of Sciences 116(52), 26555–26563. doi: 10.1073/pnas.1911837116.

Wang, K., Johnstone, M.A., Xin, C., Song, S., Padilla, S., Vranka, J.A., et al. (2017a). Estimating Human Trabecular Meshwork Stiffness by Numerical Modeling and Advanced OCT Imaging. Investigative Ophthalmology & Visual Science 58(11), 4809–4817. doi: 10.1167/iovs.17-22175.

Wang, K., Read, A.T., Sulchek, T., and Ethier, C.R. (2017b). Trabecular meshwork stiffness in glaucoma. Exp Eye Res 158, 3–12. doi: 10.1016/j.exer.2016.07.011.

Yemanyi, F., and Raghunathan, V. (2020). Lysophosphatidic Acid and IL-6 Trans-signaling Interact via YAP/TAZ and STAT3 Signaling Pathways in Human Trabecular Meshwork Cells. Invest Ophthalmol Vis Sci 61(13), 29. doi: 10.1167/iovs.61.13.29.

Yemanyi, F., Vranka, J., and Raghunathan, V.K. (2020a). Crosslinked Extracellular Matrix Stiffens Human Trabecular Meshwork Cells Via Dysregulating beta-catenin and YAP/TAZ Signaling Pathways. Invest Ophthalmol Vis Sci 61(10), 41. doi: 10.1167/iovs.61.10.41.

Yemanyi, F., Vranka, J., and Raghunathan, V.K. (2020b). Crosslinked Extracellular Matrix Stiffens Human Trabecular Meshwork Cells Via Dysregulating β-catenin and YAP/TAZ Signaling Pathways. Investigative Ophthalmology & Visual Science 61(10), 41–41. doi: 10.1167/iovs.61.10.41.

Zhang, Y., Conrad, A.H., and Conrad, G.W. (2011). Effects of ultraviolet-A and riboflavin on the interaction of collagen and proteoglycans during corneal cross-linking. J Biol Chem 286(15), 13011–13022. doi: 10.1074/jbc.M110.169813.

Zhang, Y.E. (2009). Non-Smad pathways in TGF-β signaling. Cell Research 19(1), 128–139. doi: 10.1038/cr.2008.328.

Zhao, B., Wei, X., Li, W., Udan, R.S., Yang, Q., Kim, J., et al. (2007). Inactivation of YAP oncoprotein by the Hippo pathway is involved in cell contact inhibition and tissue growth control. Genes Dev 21(21), 2747–2761. doi: 10.1101/gad.1602907.

