## Supplementary Material for "Extracellular matrix stiffness and TGFβ2 regulate YAP/TAZ activity in human trabecular meshwork cells"

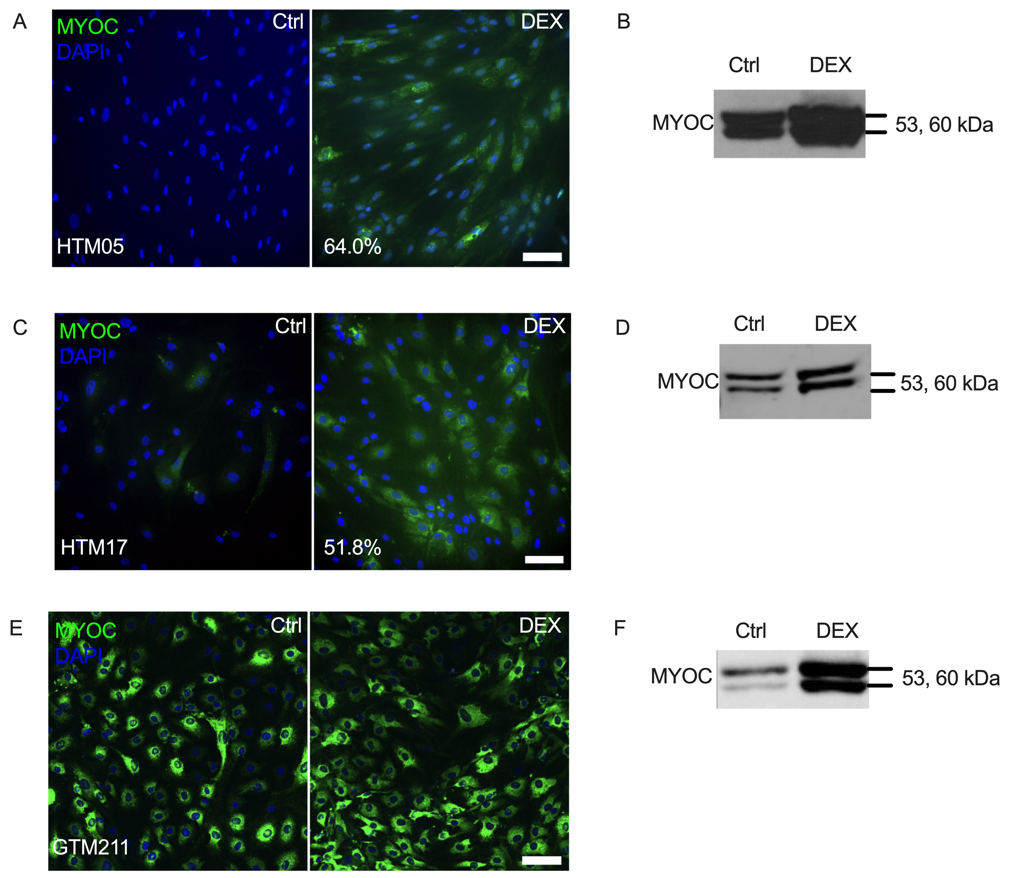

**Suppl. Fig. 1. HTM05, HTM17 and GTM211 cell characterization.** (A and C) Representative fluorescence micrographs of intracellular MYOC at 7 d (MYOC = green; DAPI = blue). Scale bar, 100 μm. (B and D) Immunoblot of secreted MYOC at 7 d.

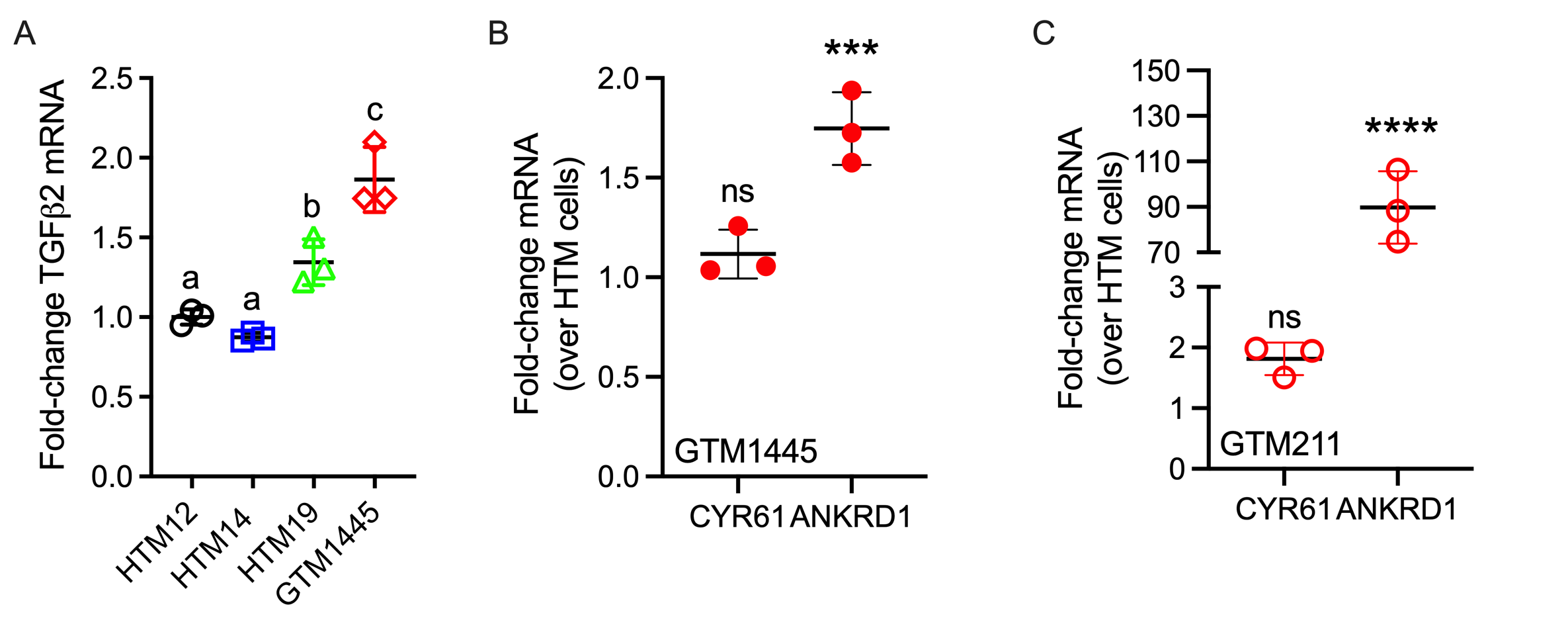

**Suppl. Fig. 2. Expression of TGFβ2, CRY61 and ANKRD1 mRNA in GTM cells compared to normal HTM cells. (A)** mRNA fold-change of TGFβ2 by qRT-PCR (N = 3 replicates/cell strain). The bars and error bars indicate Mean ± SD. Significance was determined by one-way ANOVA using multiple comparisons tests (shared significance indicator letters represent non-significant difference (p>0.05), distinct letters represent significant difference (p<0.05)).

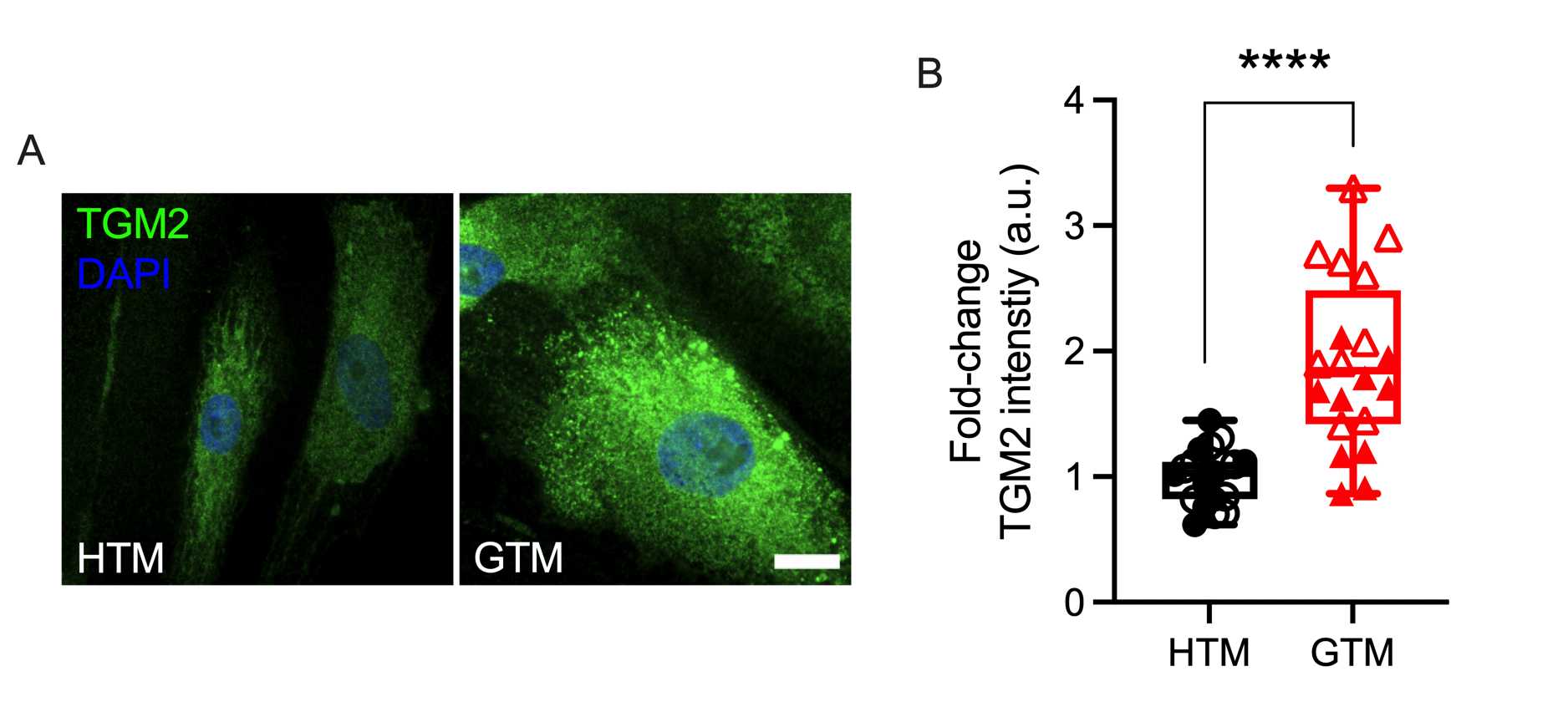

**Suppl. Fig. 3. Upregulated TGM2 expression in GTM cells compared to normal HTM cells.** (A) Representative fluorescence micrographs of TGM2 in HTM and GTM cells (TGM2 = green; DAPI = blue). Scale bar, 20 μm. (B) Analysis of TGM2 intensity in HTM and GTM cells (N = 20 images per group from 2 HTM/GTM cell strains with 6 replicates). Open and closed symbols represent different cell strains. The box and whisker plots represent median values (horizontal bars), 25th to 75th percentiles (box edges) and minimum to maximum values (whiskers), with all points plotted. Significance was determined by unpaired t- test (****p < 0.0001).

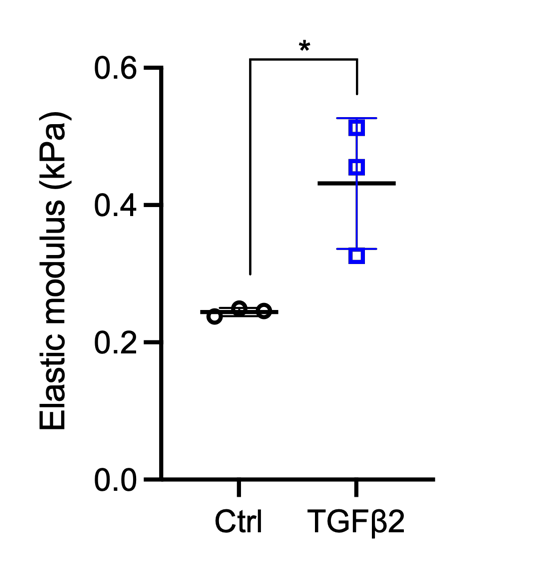

**Suppl. Fig. 4. Stiffness of HTM cell-encapsulated hydrogels.** Elastic modulus of HTM cell-encapsulated hydrogels measured by rheometer (N = 3 replicates/group). The bars and error bars indicate Mean ± SD. Significance was determined by unpaired t-test (*p<0.05).

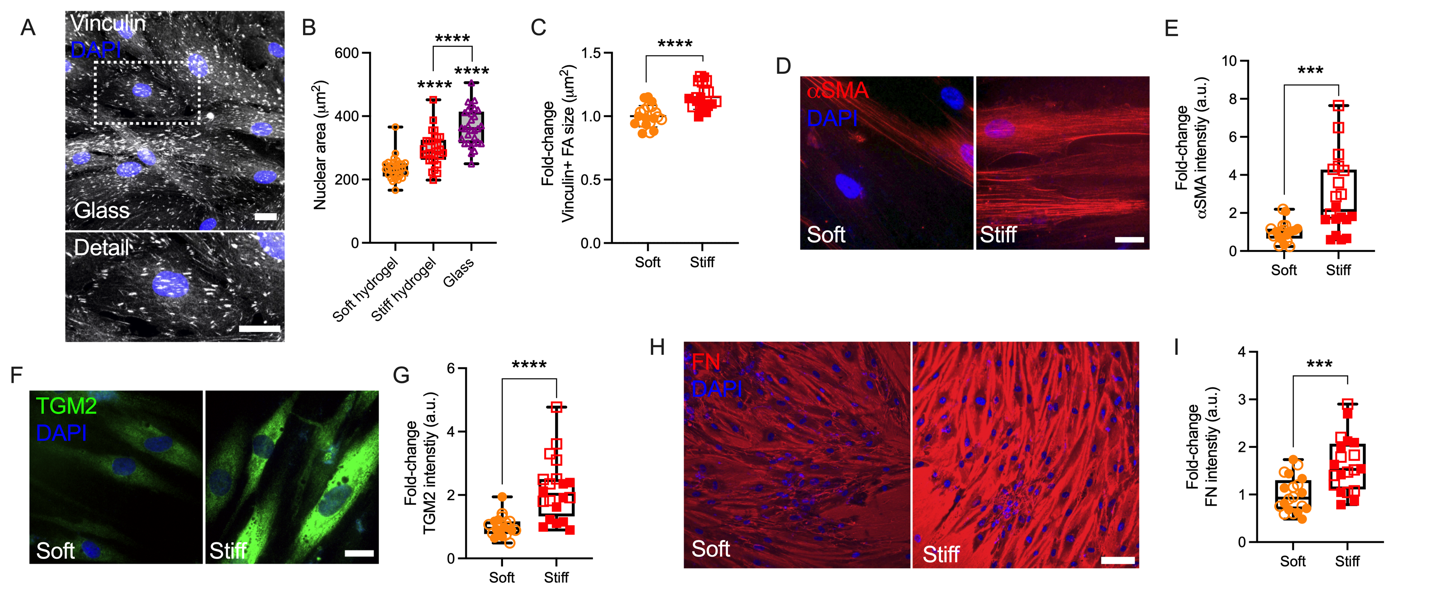

**Suppl. Fig. 5. Effects of ECM hydrogel stiffness on nuclear size, αSMA, TGM2 and FN expression.** (A) Representative fluorescence micrographs of vinculin in HTM cells on glass (dashed box shows region of interest in higher magnification image; vinculin = grey; DAPI = blue). Scale bar, 20 μm. (B) Nuclear area of HTM cells on soft and stiff hydrogels (N = 30 images from 2 HTM cell strains with 3-6 replicates per cell strain; more than 100 nuclei were analyzed per cell strain). (C) Analysis of size of vinculin puncta (N = 20 images from 2 HTM cell strains with 3 replicates per cell strain). (D, F and H) Representative fluorescence micrographs of αSMA, TGM2 and FN in HTM cells on soft and stiff hydrogels (αSMA/FN = red; TGM2 = green; DAPI = blue). Scale bar, 20 μm in C and E; 100 μm in F. (E, G and I) Analysis of αSMA, TGM2 and FN intensity (N = 20 images from 2 HTM cell strains with 3 replicates per cell strain). Open and closed symbols represent different cell strains. The bars and error bars indicate Mean ± SD. The box and whisker plots represent median values (horizontal bars), 25th to 75th percentiles (box edges) and minimum to maximum values (whiskers), with all points plotted. Significance was determined by unpaired t-test (D, F and H) or one-way ANOVA (B) using multiple comparisons tests (****p<0.0001).

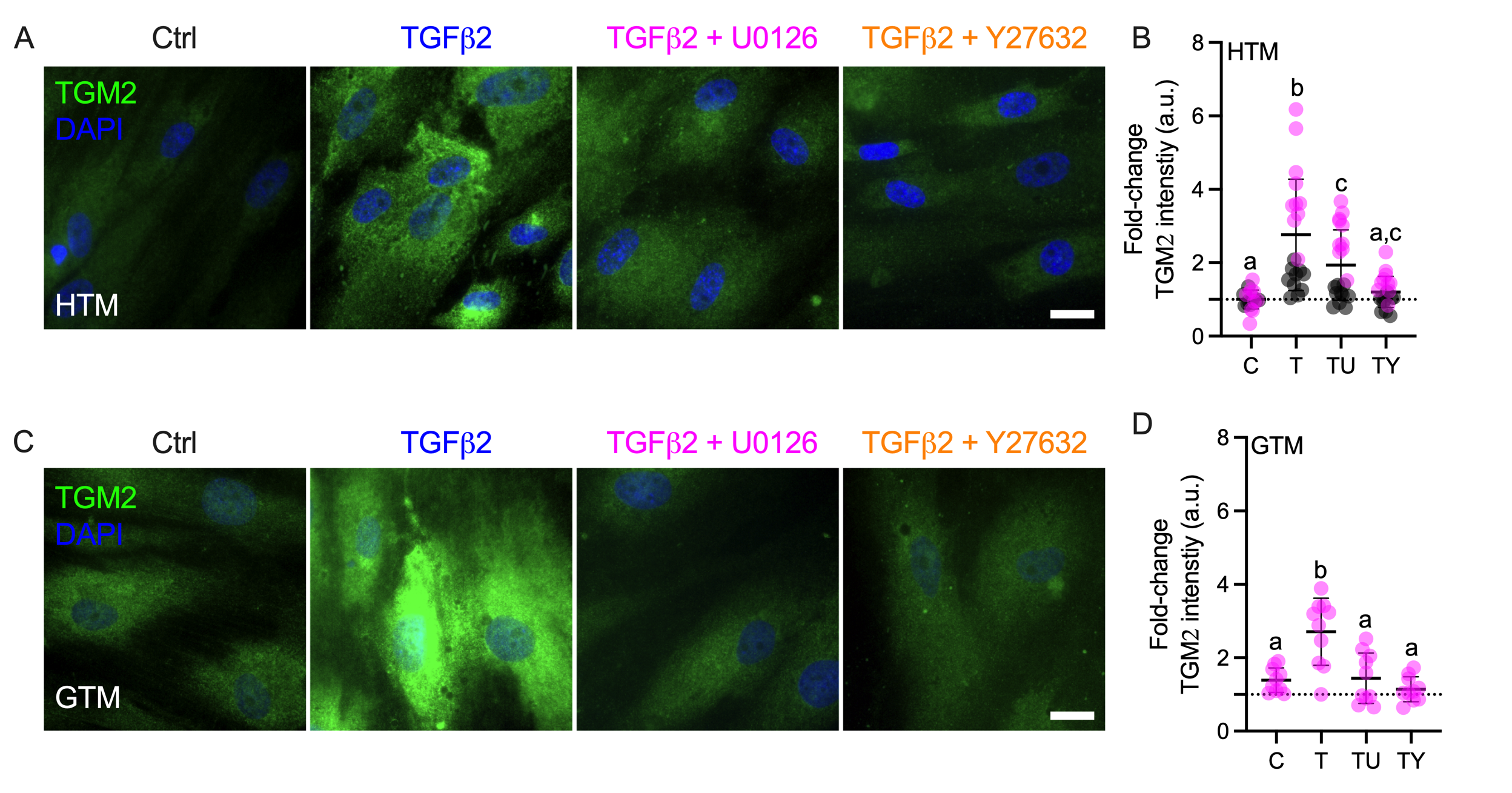

**Suppl. Fig. 6. Effects of TGFβ2 in absence or presence of ERK or ROCK inhibition on TGM2 expression.** (A and C) Representative fluorescence micrographs of TGM2 in HTM or GTM cells on soft hydrogels subjected to control, TGFβ2 (2.5 ng/mL), TGFβ2 + U0126 (10 µM), TGFβ2 + Y27632 (10 µM) at 3 d (TGM2 = green; DAPI = blue). Scale bar, 20 μm. (B and D) Analysis of TGM2 intensity in HTM and GTM cells (N = 20 images from 2 HTM cell strains with 3 replicates per cell strain; N = 10 images from one GTM cell strain with 3 replicates). Symbols with different colors represent different cell strains; dotted line shows HTM cells control value for reference. The bars and error bars indicate Mean ± SD. Significance was determined by one-way ANOVA using multiple comparisons tests (shared significance indicator letters represent non-significant difference (p>0.05), distinct letters represent significant difference (p<0.05)).

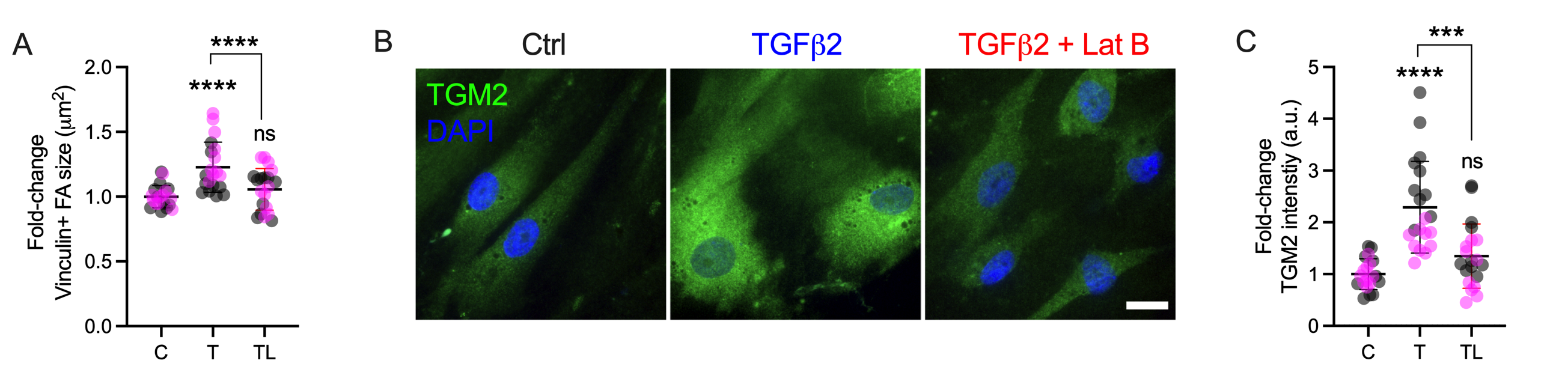

**Suppl. Fig. 7. Lat B reduces TGM2 expression in HTM cells.** (A) Analysis of size of vinculin puncta (N = 20 images from 2 HTM cell strains with 3 replicates per cell strain). (B) Representative fluorescence micrographs of TGM2 in HTM cells on soft hydrogels subjected to control, TGFβ2 (2.5 ng/mL), TGFβ2 + Lat B (2 µM) at 3 d (TGM2 = green; DAPI = blue). Scale bar, 20 μm. (C) Analysis of TGM2 intensity (N = 20 images per group from 2 HTM cell strains with 3 replicates per HTM cell strain). Symbols with different colors represent different cell strains. The bars and error bars indicate Mean ± SD. Significance was determined by one-way using multiple comparisons tests (****p < 0.0001).

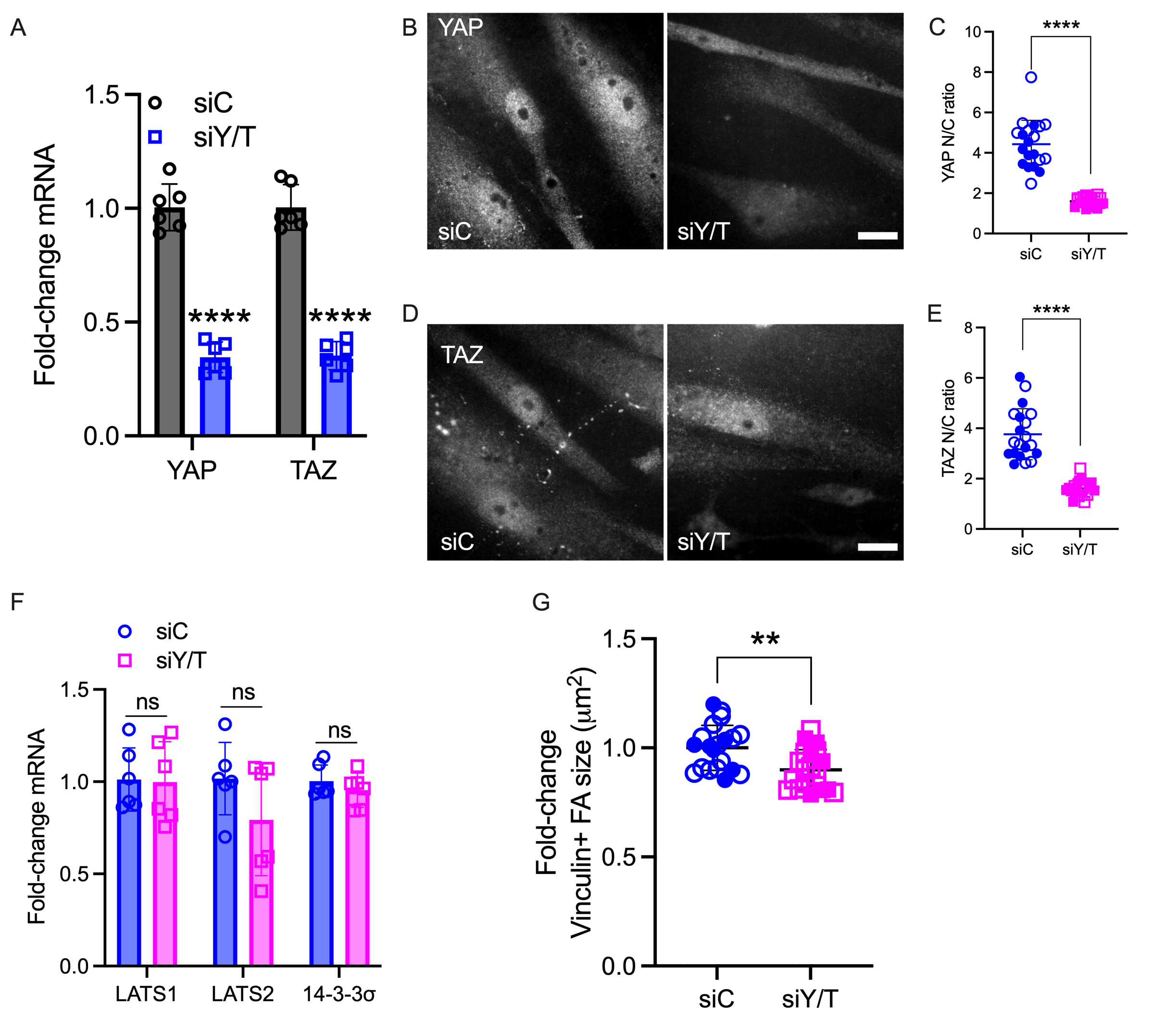

**Suppl. Fig. 8. YAP/TAZ depletion using siRNA.** (A) mRNA fold-change of YAP and TAZ in HTM cells on stiff hydrogels subjected to siControl or siYAP/TAZ knockdown by qRT-PCR. The mRNA levels were normalized to the levels of GAPDH mRNA 48h post transfection (N = 6 replicates from 2 HTM cell strain). (B and D) Representative fluorescence micrographs of YAP/TAZ in HTM cells on stiff hydrogels subjected to siControl or siYAP/TAZ (YAP/TAZ = grey). Scale bar, 20 μm. (C and E) Analysis of YAP/TAZ nuclear/cytoplasmic ratio (N = 20 images from 2 HTM cell strains with 3 replicates per cell strain). (F) mRNA fold-change of LATS1/2 and 14-3-3σ in HTM cells on stiff hydrogels subjected to siControl or siYAP/TAZ by qRT-PCR. The mRNA levels were normalized to the levels of GAPDH mRNA 48h post transfection (N = 6 replicates from 2 HTM cell strain). (G) Analysis of size of vinculin puncta (N = 20 images from 2 HTM cell strains with 3 replicates per cell strain). Open and closed symbols represent different cell strains. The bars and error bars indicate Mean ± SD; Significance was determined by unpaired t-test (D and G) and two-way ANOVA (A and F) using multiple comparisons tests (****p < 0.0001).

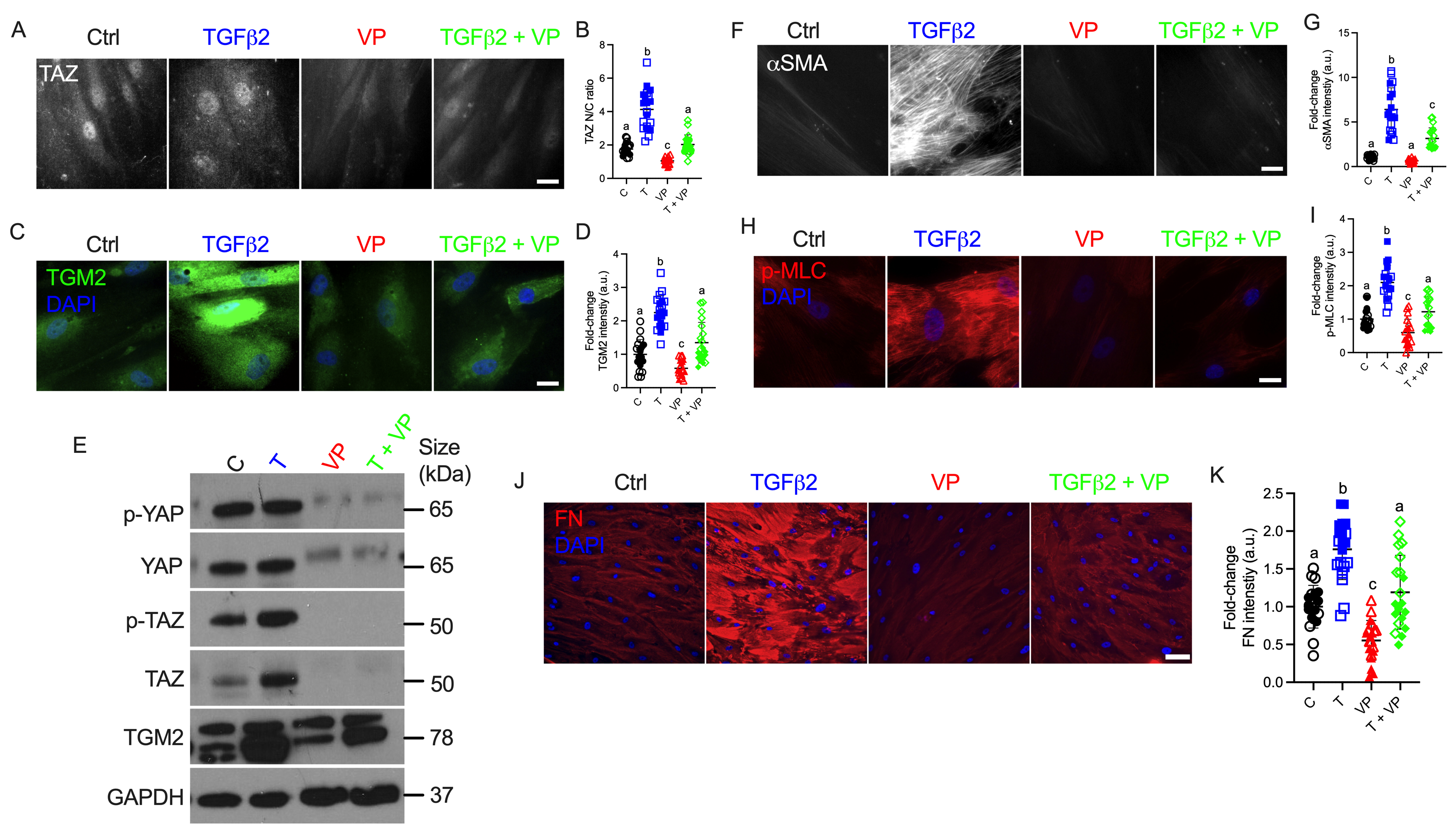

**Suppl. Fig. 9. Inhibition of YAP/TAZ-TEAD interaction with VP decreases nuclear TAZ, TGM2, cell contractile properties and ECM remodeling in HTM cells.** (A) Representative fluorescence micrographs of TAZ in HTM cells on soft hydrogel subjected to control, TGFβ2 (2.5 ng/mL), VP (0.5 µM), TGFβ2 + VP at 3 d (YAP/TAZ = red; DAPI = blue). Scale bar, 20 μm. (B) Analysis of TAZ nuclear/cytoplasmic ratio (N = 20 images from 2 HTM cell strains with 3 replicates per cell strain). (C) Representative fluorescence micrographs of TGM2 in HTM cells on soft hydrogel subjected the different treatments (TGM2 = green; DAPI = blue). Scale bar, 20 μm. (D) Analysis of TGM2 intensity (N = 20 images from 2 HTM cell strains with 3 replicates per cell strain). (E) Immunoblot of p-YAP, total YAP, p-TAZ, total TAZ and TGM2. (F, H and J) Representative fluorescence micrographs of αSMA, p-MLC and FN in HTM cells on soft hydrogel subjected to the different treatments (αSMA = grey; FN/p-MLC = red; DAPI = blue). Scale bar, 20 μm in F and H; 100 μm in J. (G, I and K) Analysis of αSMA, p-MLC and FN (N = 20 images from 2 HTM cell strains with 3 replicates per cell strain). Open and closed symbols represent different cell strains. The bars and error bars indicate Mean ± SD; Significance was determined by one-way ANOVA using multiple comparisons tests (shared significance indicator letters represent non-significant difference (p>0.05), distinct letters represent significant difference (p<0.05)).

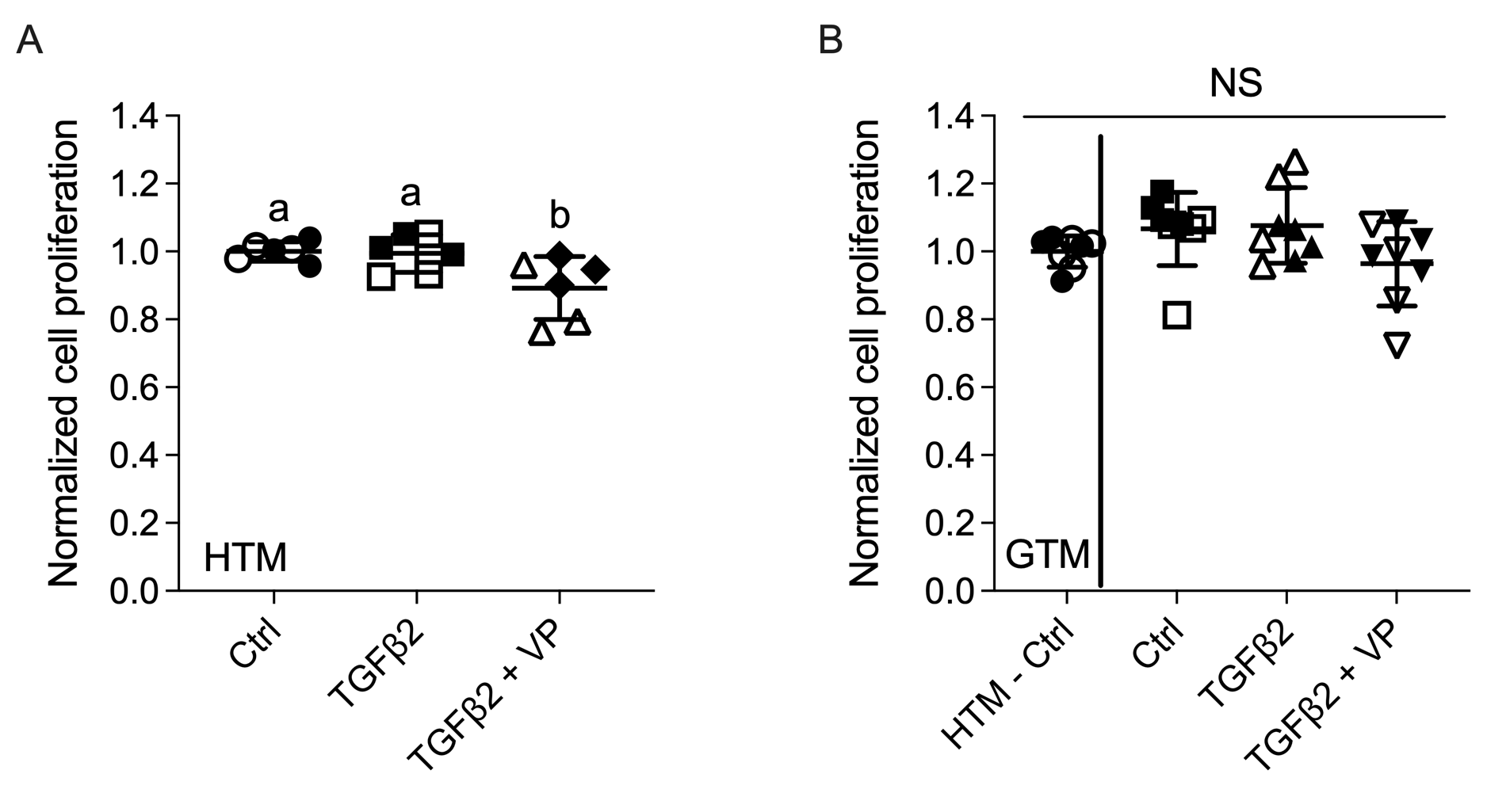

**Suppl. Fig. 10. HTM/GTM cell viability affected by YAP/TAZ inhibition.** Cell proliferation quantification of HTM hydrogels (A) and GTM hydrogels (B) subjected to control, TGFβ2 (2.5 ng/ml), TGFβ2 + VP (0.5 µM) at 5 d (N = 6-7 hydrogels per group from 2 HTM/GTM cell strains with 3-4 replicates per HTM/GTM cell strain). Open and closed symbols represent different cell strains. The bars and error bars indicate Mean ± SD; Significance was determined by one-way using multiple comparisons tests (shared significance indicator letters represent non-significant difference (p>0.05), distinct letters represent significant difference (p<0.05)).

**Table 1**

| ID | Sex | Age |
| --- | --- | --- |
| HTM05 | Male | 57 |
| HTM12 | Female | 60 |
| HTM14 | Female | 50 |
| HTM17 | Male | 33 |
| HTM19 | Male | 34 |
| GTM211* | Female | 75 |
| GTM1445* | Female | 81 |
| **obtained from W.D.S. at Duke University* | | |

**Table 2**

| **Target** | **Catalog no.** | **Company** | **Dilution WB** | **Dilution ICC** |
| --- | --- | --- | --- | --- |
| anti-p-YAP | 4911S | Cell Signaling Technology | 1:1000 |  |
| anti-p-TAZ | 59971 | Cell Signaling Technology | 1:1000 |  |
| anti-YAP | 14074S | Cell Signaling Technology | 1:1000 | 1:200 |
| anti-TAZ | 4883S | Cell Signaling Technology | 1:1000 | 1:200 |
| anti-YAP | sc-376830 | Santa Cruz Biotechnology | 1:200 |  |
| anti-TAZ | sc-293183 | Santa Cruz Biotechnology | 1:200 |  |
| anti-14-3-3𝛔 | 8312 | Cell Signaling Technology | 1:1000 |  |
| anti-TGM2 | ab421 | Cell Signaling Technology | 1:1000 | 1:400 |
| anti-GAPDH | G9545 | Sigma | 1:80000 |  |
| anti-Vinculin | ab129002 | Abcam |  | 1:200 |
| anti-p-MLC | 3675 | Cell Signaling Technology |  | 1:200 |
| anti-MYOC | MABN866 | Sigma | 1:2000 |  |
| anti-MYOC | ab41552 | Abcam |  | 1:200 |
| anti-Fibronectin | ab45688 | Abcam |  | 1:500 |
| anti-αSMA | C6198 | Sigma |  | 1:400 |
| Phalloidin | 12935S | Cell Signaling Technology |  | 1:500 |
| anti-Rabbit HRP | ab6721 | Abcam | 1:50000 |  |
| IRDye® 680RD Goat anti-Rabbit IgG | 926-68071 | LI-COR | 1:15000 |  |
| IRDye® 800CW Goat anti-Mouse IgG | 926-32210 | LI-COR | 1:15000 |  |
| Alexa Fluor® 488-conjugated anti-Rabbit IgG | A27034 | Invitrogen |  | 1:500 |
| Alexa Fluor® 584-conjugated anti-Rabbit IgG | ab150080 | Abcam |  | 1:500 |
| Alexa Fluor® 488-conjugated anti-Mouse IgG | A21203 | Invitrogen |  | 1:500 |

**Table 3**

| **Target** | **Forward** | **Reverse** |
| --- | --- | --- |
| GAPDH | GTCTCCTCTGACTTCAACAGCG | ACCACCCTGTTGCTGTAGCCAA |
| YAP | AGCCAGTTGCAGTTTTCAGG | AACAGCAGCAATGGACAAGG |
| TAZ | CATGGCAGTATCCCAGCCAA | AGCGCATTGGGCATACTCAT |
| CTGF | ATGTGCATTCTCCAGCCATC | TTCACTTGCCACAAGCTGTC |
| TGM 2 | TCAACTGCAACGATGACCAGG | TGTTCTGGTCATGGGCCGAG |
| CYR61 | GAGTGGGTCTGTGACGAGGAT | GGTTGTATAGGATGCGAGGCT |
| ANKRD1 | AGTAGAGGAACTGGTCACTGG | TGGGCTAGAAGTGTCTTCAGAT |
| 14-3-3𝛔 | TATAAGAACGTGGTGGGCGG | CCTCCTTGATGAGGTGGCTG |
| FN | GTCTTGTGTCCTGATCGTTG | AGGCTGGATGATGGTAGATTG |
| a-SMA | GGCATCATCACCAACTGGGA | CAGGGTGGGATGCTCTTCAG |
| LATS1 | CTCTGCACTGGCTTCAGATG | TCCGCTCTAATGGCTTCAGT |
| LATS2 | ACATTCACTGGTGGGGACTC | GTGGGAGTAGGTGCCAAAAA |

**Captions for videos:**

**Video 1**. 3D re-construction of confocal z-stacks of F-actin in HTM cells cultured in soft hydrogels subjected to control (turn around x-axis).

**Video 2**. 3D re-construction of confocal z-stacks of F-actin in HTM cells cultured in soft hydrogels subjected to TGFβ2 (2.5 ng/mL) (turn around x-axis).

**Video 3**. Representative 3D re-construction of confocal z-stacks of F-actin in HTM cells cultured in soft hydrogels subjected to VP (0.5 μM) (turn around x-axis).

**Video 4**. Representative 3D re-construction of confocal z-stacks of F-actin in HTM cells cultured in soft hydrogels subjected to TGFβ2 + VP (turn around x-axis).
